# Rad52 mediates class-switch DNA recombination to IgD

**DOI:** 10.1101/2021.06.28.450246

**Authors:** Yijang Xu, Hang Zhou, Ginell Post, Hong Zan, Paolo Casali

**Affiliations:** Department of Microbiology, Immunology & Molecular Genetics, University of Texas Long School of Medicine; Department of Pathology, University of Arkansas School of Medicine, Little Rock, AR 72205, USA; Department of Medicine, University of Texas Long School of Medicine, UT Health Science Center, San Antonio, TX 78229, USA

## Abstract

While the biology of IgD begins to be better understood, the mechanism of expression of this phylogenetically old and highly conserved Ig remains unknown. In B cells, IgD is expressed together with IgM as transmembrane receptor for antigen through alternative splicing of long primary *V_H_DJ_H_-Cμ-s-m-Cδ-s-m* RNAs, which also underpin secreted (s)IgD. IgD is also expressed through class switch DNA recombination (CSR), as initiated by AID-mediated double-strand DNA breaks (DSBs) in Sμ and σδ, and resolution of such DSBs by a still unknown mechanism. This synapses Sμ with σδ region DSB resected ends leading to insertion of extensive S-S junction microhomologies, unlike Ku70/Ku86-dependent NHEJ which resolves DSB blunt ends in CSR to IgG, IgA and IgE with little or no microhomologies. Our previous demonstration of a novel role of Rad52 in a Ku70/Ku86-independent “short-range” microhomology-mediated synapsis of intra-Sμ region DSBs led us to hypothesize that this homologous recombination DNA annealing factor is also involved in short-range microhomology-mediated alternative endjoining (A-EJ) recombination of Sμ with σδ. We found that induction of IgD CSR by selected stimuli downregulated Zfp318 (the suppressor of *Cμ-s-m* transcription termination), promoted Rad52 phosphorylation and Rad52 recruitment to Sμ and σδ, leading to Sμ-σδ recombination with extensive microhomologies, *V_H_DJ_H_-Cδs* transcription and sustained IgD secretion. Rad52 ablation in mouse *Rad52^−/−^* B cells aborted IgD CSR *in vitro* and *in vivo* and dampened the specific IgD antibody response to OVA. Further, Rad52 knockdown in human B cells virtually abrogated IgD CSR. Finally, Rad52 phosphorylation was associated with high levels of IgD CSR and anti-nuclear IgD autoantibodies in lupus-prone mice and lupus patients. Thus, Rad52 effects CSR to IgD through microhomology-mediated A-EJ and in concert with Zfp318 modulation. This is a previously unrecognized, critical and dedicated role of Rad52 in mammalian DNA repair that provides a mechanistic underpinning to CSR A-EJ.

---

IgD has been an enigmatic antibody class for many years, despite being evolutionarily ancient and highly conserved across species^1–6^. As primordial as IgM, IgD appeared in cartilaginous fishes, amphibians and occurs in fishes, rodents, cattle and humans^2, 7^. As an example, in *Xenopus*, the Igδ exon cluster is in the same position, immediately 3′ of the Igμ locus, as it exists in mammals^7^. In mice and humans, IgD is expressed primarily as a transmembrane IgD receptor together with IgM with identical antigen specificity on naïve mature B cells in the form of BCR. IgD also exists as a secreted antibody. In humans, circulating IgD occurs at concentrations up to more than two-thousand folds greater than IgE (10-250 μg/ml vs. ∼0.1 μg/ml), the rarest peripheral blood Ig class. IgD are secreted by IgM^−^IgD^+^ plasmablasts and plasma cells differentiated from B cells in lymphoepithelial organs in aerodigestive mucosae, including palatine and pharyngeal tonsils. IgM^−^IgD^+^B cells and plasma cells can also be found in the lachrymal, salivary and mammary glands^3^. In addition to existing as free molecule, IgD can occurs on the surface of innate effector cells, including basophils, mast cells and monocytes^1, 8, 9^. IgD bound to these cells would enhance immune surveillance and exert proinflammatory and antimicrobial effects^1, 8, 9^. These include triggering basophils to secret IL-4, IL-5 and IL-13 upon antigen engagement or attenuating basophil or mast cell allergic degranulation induced by IgE co-engagement^1^. Thus, IgD would contribute to mucosal homeostasis by endowing effector cells with reactivity to microbial commensals and pathogens^5, 6^.

Identifying the stimuli and molecular mechanisms that underpin IgD expression is important to understand the regulation of IgD secretion throughout the body. The immediately proximal location and unique integration of Cδ and Cµ gene loci in the same transcriptional unit allow these two Ig isotypes to be coordinately regulated in transcription^10, 11^. In naive mature B cells, (membrane) mIgM and mIgD are co-expressed by alternative splicing of long primary transcripts consisting of rearranged V_H_DJ_H_ exons and downstream Cµ and Cδ exons (*V_H_DJ_H_-Cµ-s-m-Cδ-s-m*). Alternative splicing of the same long primary *V_H_DJ_H_-Cµ-s-m-Cδ-s-m* transcripts also leads to expression of (secreted) sIgM and sIgD^2, 8^. Transcription of long primary *V_H_DJ_H_-Cµ-s-m-Cδ-s-m* RNA requires the zinc finger ZFP318 repressor of transcriptional termination, which obliterates the effect of the transcriptional termination sites (TTS) intercalated between the Cμ and Cδ exon clusters^10, 11^ (Fig. 1a). IgD can also be expressed through class switch DNA recombination (CSR), by which IgM^+^IgD^+^B cells juxtapose *V_H_DJ_H_* DNA from the Cµ (IgM) to the Cδ (IgD) exons cluster, giving rise to *V_H_DJ_H_-Cδm* RNA transcripts and IgM^−^IgD^+^B cells^1, 5, 8, 9, 12^ (Fig. 1b). In human and mouse nasopharyngeal and intestinal lymphoid tissues, a significant proportion of mucosal B cells class-switch to IgM^−^IgD^+^B cells, which subsequentially differentiate to plasmablasts and plasma cells^1, 3, 5, 6^. Generally, CSR to IgD (Cδ) is a less frequent event than CSR to IgG (Cγ), IgA (Cα) or IgE (Cε), perhaps a reflection among other factors of the peculiar structure of the pseudo-switch σδ region lying immediately upstream of Cδ exons. Compared to the canonical Sμ, S*γ*, S*α* and S*ε* regions lying 5’ of the respective Igμ, Ig*γ*, Ig*α* and Ig*ε* loci, σδ is shorter and contain differing motifs of nucleotide (nt) repeats^2, 5, 8, 13, 14^. These would provide an unconventional substrate for AID-mediated insertion of DSBs, possibly more prone to end-resection and generation of single-strand overhangs for Sμ-σδ recombination, which leads to expression of post-recombination *V_H_DJ_H_-Cδ* RNA transcripts^2, 8, 13–15^.

**Fig. 1.**
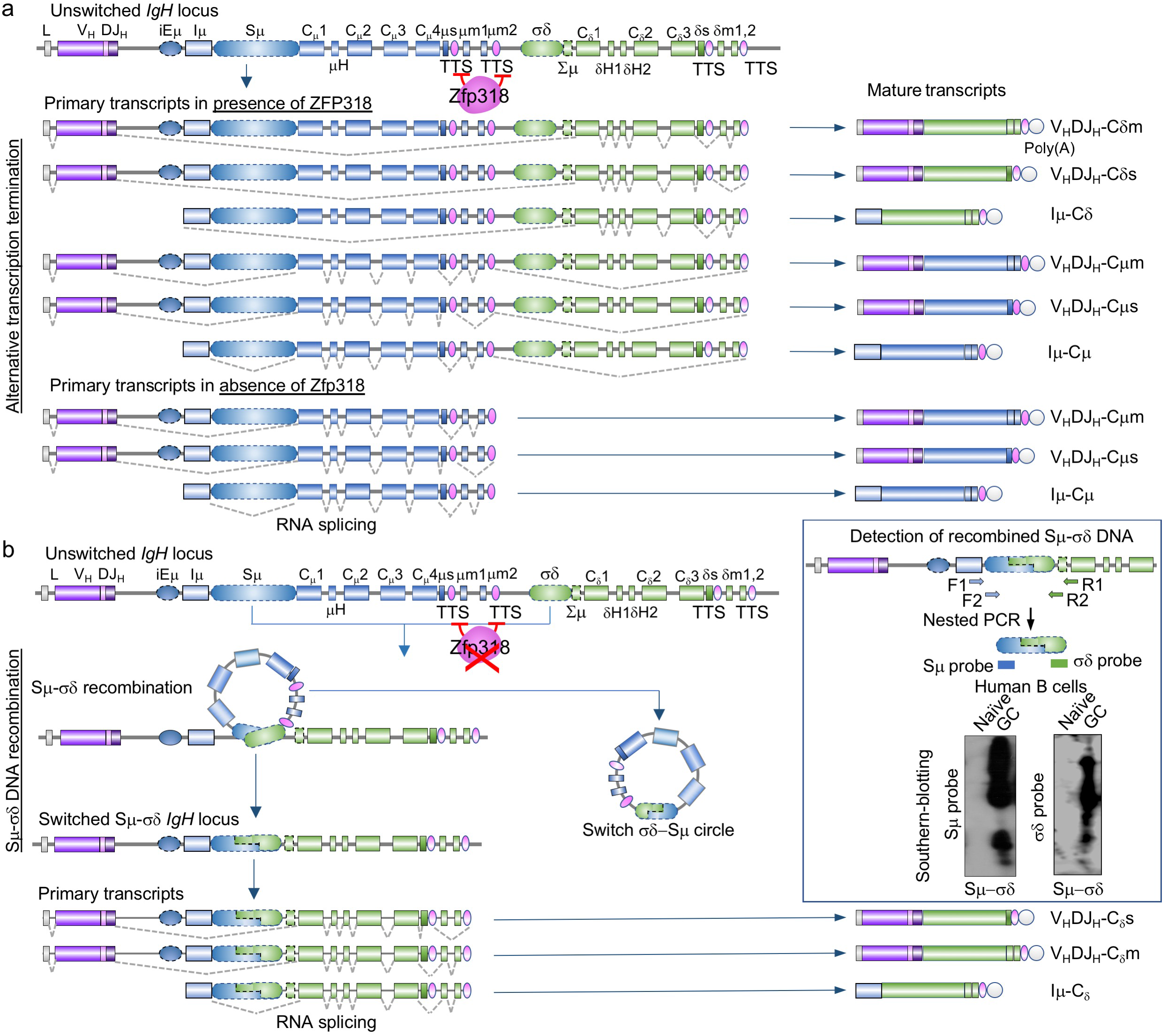
Expression of cell surface and secreted IgD and IgM, as well as Iμ-Cδ transcripts by alternative splicing, alternative transcription termination and CSR. **a**, Schematics of alternative splicing and alternative transcription termination for expression of membrane and secreted IgM and IgD, as well as germline *Iμ-Cμ* and *Iμ-Cδ* transcripts in B cells. In the presence of Zfp318, which represses the transcription termination sites (TTS) of the Cμ gene, mature B cells constitutively transcribe long primary *V_H_DJ_H_-Cμ-Cδs–m* transcripts initiated by the V_H_ promoter. These long primary transcripts undergo alternative splicing which removes intronic regions, leading to dual expression of mature *V_H_DJ_H_-Cμs* and *V_H_DJ_H_-Cδm* transcripts encoding IgM and IgD. In the absence of Zfp318, transcription stops at Cμ TTS, resulting in a shorter primary transcript, which does not contain Cδ exons, and lead to expression of a mature *V_H_DJ_H_-Cμ−s–m* transcript only. Mature B cells also transcribe *Iμ, Cμ,* and *Cδ* regions under control of the Iμ promoter. When Zfp318 is present, unswitched mature B cells constitutively transcribe long primary *Iμ-Cμ−s−m-Cδ–s−m* transcripts, which undergo alternative splicing to removes intronic regions, leading to dual expression of germline *Iμ-C*μ and *Iμ-Cδ* transcripts. In the absence of Zfp318, *Iμ* promoter-initiated transcription stops at Cμ TTS, and only germline Iμ-Cμ transcripts are expressed. **b,** Expression of membrane and secreted IgD, and *Iμ-Cδ* transcripts by CSR. Schematic representation of CSR from IgM to IgD. The Sμ region recombines with the σδ region and loops out the intervening DNA, which forms a switch circle. The recombined DNA is transcribed leading to expression of *V_H_DJ_H_-Cδ−s–m* and *Iμ-Cδ* transcripts, initiated by the V_H_ and Iμ promoters, respectively. In this case, *Iμ-Cδ* transcripts are generated as post-recombination transcripts. Graphics depict portion of the *IgH* locus and the resulting primary and mature transcripts. The inset depicts a schematic representation of the detection of Sμ*−σδ*junctional DNA (CSR to IgD) by nested PCR amplification followed by Southern-blotting using specific Sμ and σδ probes (Southern-blotting of amplified recombined Sμ*−σδ* DNA from human naïve and germinal center B cells). In many cases the amplified Sμ-σδ DNA was sequenced for further analysis of junctional sequence well as dentification and census of mutations. iEμ, *IgH* intronic enhancer; Iμ, intervening μ exon; μm, exon encoding the transmembrane region of IgM; δm, exon encoding the secretory piece of IgM; σδ, noncanonical switch-like region 5′ to Cδ; δs, exon encoding the secretory region of IgD; Cδm, exon encoding the transmembrane region of IgD. Dotted gray lines show splicing configurations of primary transcripts to yield secreted and transmembrane forms of IgM and IgD.

Unlike CSR to IgG, IgA and IgE, the mechanism of CSR to IgD remains unknown. Recombination involving Sμ DSB ends with DSB ends in downstream Sμ, S*γ*, S*α* or S*ε* region is effected by non-homologous end-joining (NHEJ), one of the two major DNA DSB repair pathways, the other being homologous recombination (HR)^16, 17^. HR accurately repairs resected (staggered) DSB ends using a sister chromatid as a homologous single-strand template during cell cycle S-G2. It critically effects error-free DSB repair in somatic cells and helps orchestrate chromosome segregation in meiosis. In contrast to HR, NHEJ is a homology-independent error-prone process. It synapses blunt or virtually blunt DSB ends that lack substantial joining complementarity to form “direct” junctions, predominantly in G1 but also throughout the whole cell cycle^16^. NHEJ requires Ku70/Ku86 and in CSR mediates efficient long-range synapses of Sμ DSB ends with S*γ*, S*α* and S*ε* DSB ends, leading to IgG, IgA and IgE^15^. The finding, however, that reduction or deletion of Ku70/Ku86 led to reduced but still substantial CSR to IgG1 and IgG3 supported the existence of an alternative CSR end-joining (A-EJ) pathway^18–20^. This, like HR, would join resected DSB ends, thereby giving rise to S-S junctions with microhomologies. Unlike HR, however, the A-EJ pathway juxtaposes DSB overhangs to be joined without using a homologous template as a guide. Rather, it utilizes differing extents of sequence complementarity (homology) between the upstream and downstream resected DSB overhangs to align the to-be DNA junctions^21^. As we have shown, HR factor Rad52 competes with Ku70/Ku86 for binding to S region DSB ends and synapses DSB ends by A-EJ through microhomology-mediated end-joining (MMEJ)^20^, as inferred from increased NHEJ-mediated IgG, IgA and IgE CSR events with even fewer S-S junction microhomologies in *Rad52^−/−^* B cells *in vivo* and *in vitro*^20^. This together with the increased CSR to IgD in B cells lacking 53BP1, which protects S regions DSB ends from resection and facilitates long-range NHEJ to IgG, IgA and IgE^22–24^, as well as other findings of ours showing reduced intra-Sμ DSB short-range rejoining in *Rad52^−/−^* B cells^20^ led us to hypothesize that by annealing to single-strand resected DSB ends, Rad52 mediates CSR to IgD through A-EJ involving short-range Sμ-σδDSB recombination.

To test the hypothesis that Rad52 synapses Sµ with σδ DSB ends for IgD CSR, we first set up to define the stimuli that consistently induce CSR to IgD in mouse and human B cells. We then used such stimuli in mouse *Rad52^−/−^*B cells and *RAD52* siRNA knockdown human B cells together with molecular genetic methods to determine the impact of Rad52 deficiency as well as Rad52 phosphorylation on Sµ-σδ DNA recombination and IgD expression. We validated our findings by analyzing specific IgD antibody and total IgD titers in mouse blood, lungs and gut, as well as recombined Sµ-σδ DNA sequences in mouse spleen, mesenteric lymph nodes (MLNs) and Peyer’s patches as well as human tonsil B cells. We adapted chromatin immunoprecipitation (ChIP) assays to analyze the recruitment of Rad52/RAD52 to the σδ region in mouse and human B cells induced to undergo CSR to IgD, in which we also analyzed regulation of *V_H_DJ_H_-Cδ* transcription. We found that different stimuli induced IgD expression by alternative splicing of long *V_H_DJ_H_-Cµ-s-m-Cδ-s-m* RNA transcripts or by *Sµ-σδ* CSR. Further, we determined the expression of IgD by CSR to be related to Zfp318-mediated repression of the TTS integrated in Cμ-Cδ loci. We also correlated Sµ-σδ CSR with IgD secretion and plasma cell differentiation. Finally, we analyzed B cell Rad52 phosphorylation in lupus patients and lupus-prone mice and correlated it with CSR to IgD involving extensive microhomologies and somatic mutations in Sµ-σδ junctional sequences, as well as the occurrence of high levels of anti-nuclear antigen IgD autoantibodies. Our findings show that Rad52 mediates CSR to IgD through microhomology-mediated A-EJ and in concert with Zfp318 modulation. This is a previously unrecognized, critical and dedicated role of Rad52 in an essential DNA repair process in mammals.

## Results

### Definition of stimuli that induce Sµ-σδ CSR in mouse and human B cells

Toward testing our hypothesis that Rad52 mediates CSR to IgD, we first determined the stimuli that induce IgM^+^IgD^+^ B cells to undergo Sµ–σδ recombination. In these B cells, mIgD and sIgD and IgM are expressed by alternative splicing of long primary *V_H_DJ_H_-Cµ-s-m-Cδ-s-m* mRNAs – the Cδ locus is located immediately downstream of the Cµ locus in the same transcriptional unit, allowing these two loci be coordinately regulated at the transcriptional level^1, 2, 4, 6^ (Fig. 1a). As CSR can be induced in a T-dependent or T-independent antibody fashion^15, 25^, we used CD40 ligand CD154 (for mouse and human B cells), TLR4 ligand LPS (mouse B cells) and TLR9 ligand CpG (human B cells) in conjunction with differing cytokines and/or BCR-cross-linking to induced CSR to IgD. Recombined *Sµ–σδ, Sµ– Sγ1, Sµ–Sγ3, Sµ–Sα* and *Sµ–Sε* DNAs were detected by specific nested PCRs followed by positive identification of amplified DNA by blotting and hybridization with specific DNA probes (Fig. 1 inset), complemented by sequencing of the junctional Sµ-σδ or Sµ-S_X_ DNA. Of all stimuli used, only LPS or CD154 plus IL-4 induced CSR to IgD in mouse B cells (Fig. 2a), and only CpG plus IL-2 and IL-21, or CD154 plus IL-4 or IL-15 and IL-21 induced CSR to IgD in human B cells. IgD CSR was also detected *in vivo* in tonsil B cells (Fig. 2b). The effectiveness of the stimuli that did not induce CSR to IgD was verified by the respective induction of the expected Sµ–S*γ*1, Sµ–S*γ*3, Sµ–S*α* or Sµ–S*ε* DNA recombination. (IgG1, IgG3, IgA or IgE) (Fig. 2a) – no CSR to IgD, IgG, IgA or IgE occurred in *Aicda*^−/−^ B cells. In all cases, CSR was further confirmed by detection of post-recombination *Iμ-Cγ1, Iμ-Cγ3, Iμ-Cα* and *Iμ-Cε* transcripts at 72 h of culture – as post-recombination Iμ-Cδ transcripts are indistinguishable from germline Iμ-Cδs-m RNA transcripts and consistent with high levels of the latter in naïve B cells, Iμ-Cδ amplification products were less abundant in class-switched IgD than naïve B cells (Figs. 1, 2c). Thus, only select stimuli induce CSR to IgD in mouse and human B cells.

**Fig. 2.**
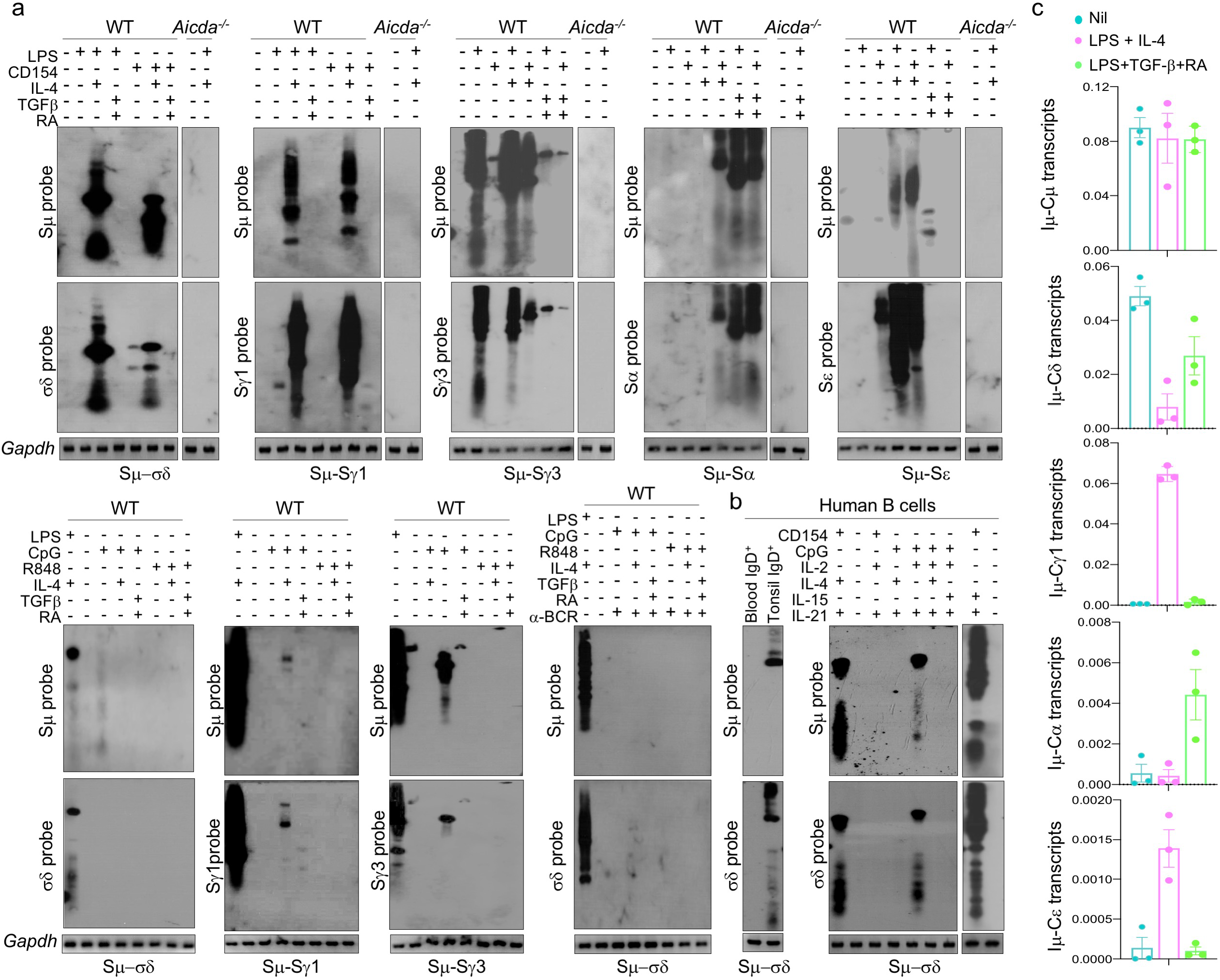
Identification of stimuli inducing CSR to IgD and Sμ-σδ junctions in mouse and human B cells. **a,** Wildtype C57BL/6 and *Aicda^−/−^* mouse naïve B cells were stimulated with nil, LPS, LPS plus IL-4, LPS plus TGF-*β* and RA, CD154, CD154 plus IL-4, CD154 plus TGF-*β* and RA, CpG, CpG plus IL-4, CpG plus TGF-*β* and RA, R848, R848 plus IL-4, R848 plus TGF-*β* and RA, or CpG plus IL-4, or R848 plus IL-4 in the presence of anti-BCR. Recombined Sμ-σδ, as well as Sμ-S*γ*1, Sμ-S*γ*3, Sμ-S*α* and Sμ-S*α* DNA were analyzed 72 or 96 h post-stimulation by nested long-range PCR using forward Iμ and reverse Cδprimers, or forward Iμ and reverse S*γ*1, S*γ*3, S*α* or S*ε* primers, respectively, followed by Southern-blotting using a specific Sμ, σδ, S*γ*1, S*γ*3, S*α* or S*ε* probe, as indicated. **b,** Recombined Sμ-σδ DNA in human tonsil IgD^+^ B cells, blood naive B cells, or blood naive B cells stimulated with CD145 or CpG plus IL-2 and IL-21, IL-4 and IL-21, or IL-2, IL-4 and IL-21 or IL-15 and Il-21, were analyzed 96 or 120 h post-stimulation by nested long-range PCR using forward Iμ and reverse Cδ primers, followed by Southern-blotting using specific human Sμ or σδ probe. Data are one representative of 3 independent experiments yielding comparable results. **c,** Germline/post-recombination Iμ-Cδtranscripts, post-recombination Iμ-C*γ*1, Iμ-C*α* and Iμ-C*ε* transcripts in wildtype C57BL/6 B cells stimulated with nil, LPS plus IL-4, or LPS plus TGF-*β* and RA, analyzed 72 h post-stimulation by qRT-PCR and normalized to *β-Actin* expression. Each dot represents data obtained with B cells from an individual mouse (*n* = 3 per group). Data are mean ± SEM.

### Sµ-σδ junctions are enriched in microhomologies and abetted by somatic mutations in mouse and human B cells

The mechanisms effecting CSR S-S synapses can leave a S-S junctional signature^20, 26^. As we previously showed, Rad52 mediates A-EJ of resected DSB ends by juxtaposing overhangs with nucleotide complementarities, thereby giving rise to Sμ-Sx DNA junctions with microhomologies^20^. Next generation sequencing of more than 100,000 recombined Sμ-Sx DNA junctions from mouse and human B cells *in vitro* and/or *in vivo* showed that Sµ–σδ junctions contained significantly more microhomologies (*p* <0.01) than Sµ–S*γ*1 or Sµ–S*α* DNA junctions (representative frequencies and lengths of microhomologies in human and mouse B cells are depicted in Fig. 3a; representative human and mouse intra-σδ and junctional Sμ-Sx sequences are depicted in Fig. 4 and Extended Data Figs. 1,2), indicating that a MMEJ^21^ synaptic process underpinned Sµ–σδ junction formation. In both human and mouse B cells, the microhomologies in Sµ–σδ junctions were significantly more extensive than those in Sµ– S*γ*1 and, to a lesser extent, Sµ–S*α*junctions (Fig. 4, Extended Data Figs. 1,2). As one example, in human tonsil B cells, 100% of analyzed Sμ-σδ junctions contained microhomologies, consisting of 2 to 13 nucleotides (mean = 6.30), while only 21% of Sµ–S*γ*1 junctions contained microhomologies, consisting of 1 to 6 nucleotides (mean = 0.72) (Fig. 3a). Interestingly, there were a few common S-S sequences shared by recombined Sµ–σδ DNA junctions in human tonsil B cells and blood naïve B cells stimulated *in vitro* by CpG plus IL-2 and IL-21, suggesting that Sµ and σδ DSB hotspots underpin Sµ–σδ in DNA recombinations. A high frequency of microhomologies was also evident in the synaptic repair process of intra-σδ DSBs, evocative of what we showed in intra-Sμ DSBs^20^. Consistent with the greatest occurrence of microhomologies in Sµ–σδ junctions, Sμ is better suited for complementary DNA single-strand annealing with σδ than S*γ*1 or, to a lesser extent, S*α* (mouse) or S*α*1 (human), based on various numbers and contexts of these DNA regions discrete motifs, such as [G_n_]AGCT repeats (Sµ, S*γ* and S*α*) or AGCTGAGCTG repeats (Sμ and σδ), as revealed by Pustell Matrix dot-plot analysis (Fig. 3b). Finally, Sµ–σδ DNA junctions were associated with somatic point-mutations. These were more frequent in the σδ area than Sμ area abetting the Sµ–σδ junction (e.g., 0.559 x 10*^−^*^2^ vs. 0.973 x 10*^−^*^2^ change/base in mouse spleen B cells *in vivo* and 1.251 x 10*^−^*^2^ vs. 1.985 x 10*^−^*^2^ change/base in mouse B cells stimulated by LPS plus IL-4 *in vitro*) (Fig. 3c). Thus, the high frequency of microhomologies in Sµ–σδ junctions supports a role of Rad52 in mediating CSR to IgD.

**Fig. 3.**
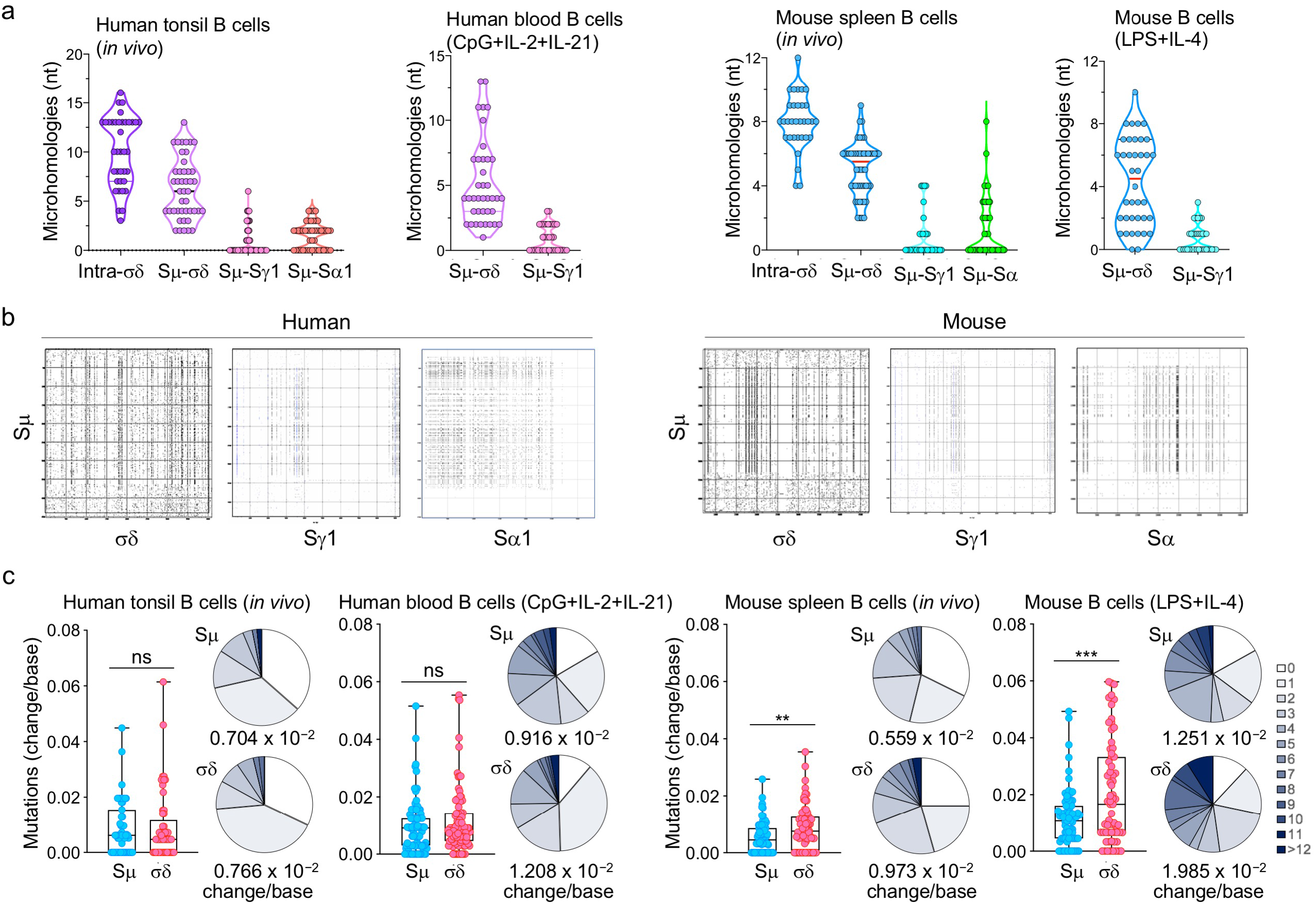
Mouse and human Sμ-σδ DNA recombination junctions contain microhomologies and somatic mutations. **a,** Amplified junctional DNAs of intra-σδ deletions, as well as Sμ-σδ, Sμ-S*γ*1 and Sμ-S*α*1 recombination from OVA-immunized C57BL/6 mouse spleen B cells, C57BL/6 mouse naïve B cells stimulated with LPS plus IL-4 and cultured for 96 h, as well as human tonsil B cells or human peripheral blood naïve B cells stimulated with CpG plus IL-2 and IL-21 and cultured for 96 or 120 h were sequenced by MiSeq. The length and numbers of nucleotide overlaps (microhomologies) in intra-σδ deletions, Sμ-σδ, Sμ-S*γ*1 and Sμ-S*α*1 junctional DNAs are shown by violin plots. Each dot represents a unique junctional sequence (*n* = 45 per group). **b,** Mouse and human Sμ and σδ regions consist of repetitive motifs, which are better suited substrates for Rad52-mediated MMEJ than those in Sμ and S*γ*1 or Sμ and S*α*. As such, they can facilitate the formation of microhomologies. Repetitive sequence elements in mouse and human Sμ, σδ, S*γ*1 and S*α* that can potentially form microhomologies were identified by Pustell Matrix dot plot using MacVector software and are depicted by small dots. Intensity of dots depicts frequency and degree of complementarity of respective sequences. **c**, Somatic point-mutations in Sμ and σδ regions abetting recombined Sμ*−σδ* DNA junctions in IgD class-switched mouse and human B cells *in vitro* and *in vivo.* Mutations were identified in a 48 to 506 nt stretch of Sμ or σδregions in unique Sμ-σδDNA recombination sequences. Each dot represents an individual sequence. Sequence data were pooled from 3 individuals in each group. In pie charts, the size of slices denotes the proportion of sequences with the same number of mutations and the grey hue denotes the number of point-mutations per sequence; below the pie charts is the overall mutation frequency (change/base). \*\**p<* 0.01, \*\*\**p <* 0.001, ns: not significant (unpaired *t* test).

**Fig. 4.**
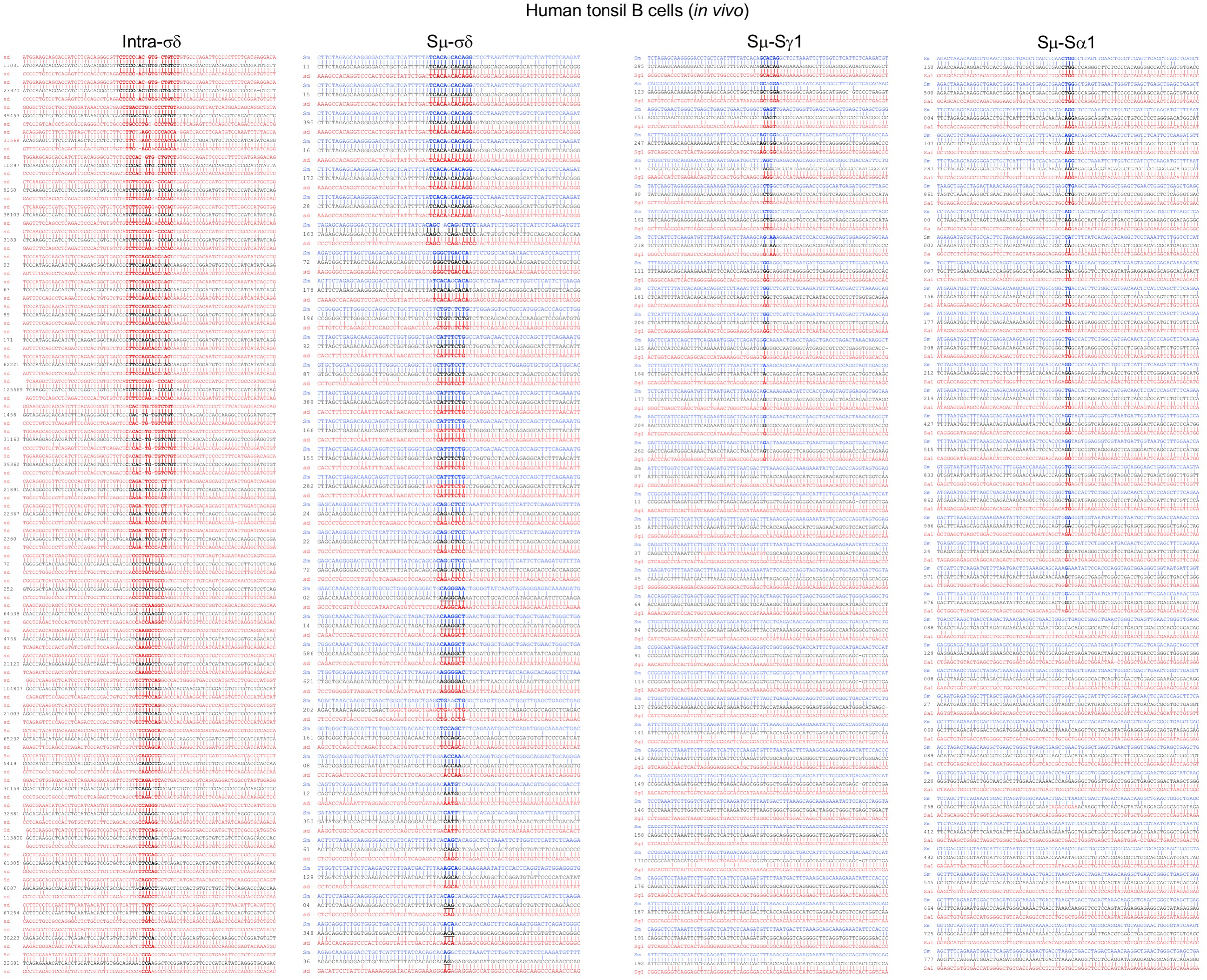
*In vivo* human Sμ-σδ DNA recombination junctions contain high frequencies of microhomologies. Amplified intra-σδ deletional, Sμ-σδ, Sμ-S*γ*1 and Sμ-S*α*1 junctional DNAs from human tonsil B cells were sequenced using MiSeq system. Thirty-two intra-σδ, Sμ-σδ, Sμ-S*γ*1 or Sμ-S*α*1 junction sequences are shown in each column. Each sequence is compared with the corresponding germline Sμ (above, blue) and σδ, S*γ*1 or S*α*1 (below, red) sequences. Microhomologies (bold) were determined by identifying the longest region at the Sμ-σδ, Sμ-S*γ*1 or Sμ-S*α*1 junction of perfect uninterrupted donor/acceptor identity or the longest overlap region at the S–S junction with no more than one mismatch on either side of the breakpoint.

### Rad52 is critically required for Sµ–σδ recombination

Having established that LPS or CD154 pus IL-4 induced CSR to IgD in mouse B cells, we used these stimuli and *Rad52^−/−^* B cells together with appropriate controls (LPS alone, LPS plus TGF-*β* and RA, CD154 or CD154 plus TGF-*β* and RA) and the same approach used in the experiments of Fig. 2 to investigate whether or not Rad52 was required for CSR to IgD. LPS plus IL-4 and CD154 plus IL-4 failed to induce Sµ–σδ recombination in *Rad52^−/−^* B cells, while either treatment efficiently induced Sμ-S*γ*1 and Sμ-S*ε* recombinations in the same *Rad52^−/−^* B cells, and Sµ–σδ recombination in *Rad52^+/+^* B cells (Fig. 5a) – in *Rad52^−/−^* B cells, LPS and LPS or CD154 plus TGF-*β* and RA induced CSR to IgG3 and IgA, respectively. As expected, CSR to IgD as well as IgG, IgA and IgE was ablated in *Aicda^−/−^* B cells. Finally, the failure of *Rad52^−/−^* B cells to undergo CSR to IgD was associated with significantly decreased secretion of IgD (Fig. 5b). Thus, Rad52 is critical for Sµ–σδ DNA recombination and seemingly important for IgD secretion.

**Fig. 5.**
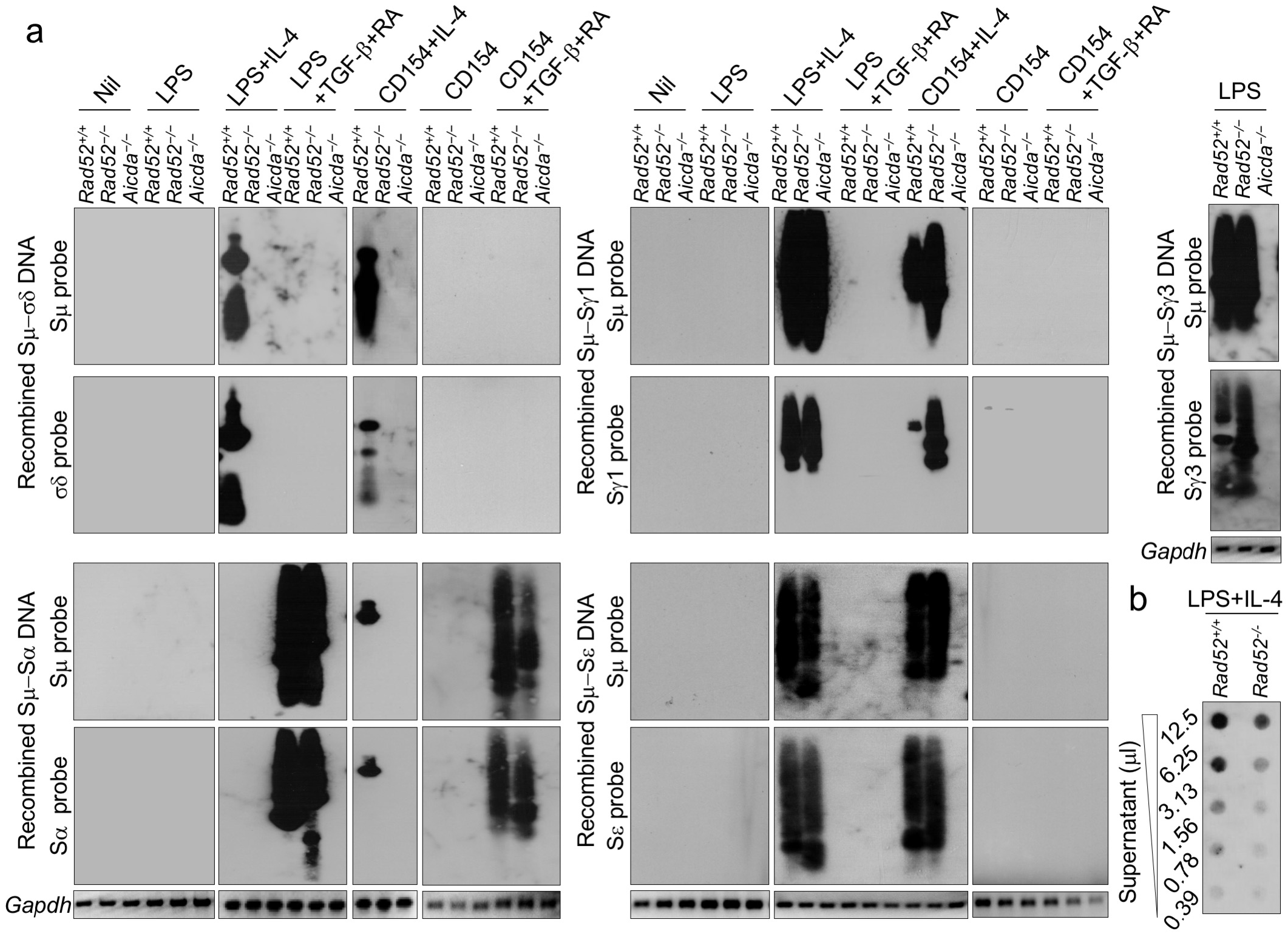
Rad52 mediates Sμ-σδ DNA recombination leading to IgD secretion. **a,** Recombined Sμ-σδ, Sμ-S*γ*1, Sμ-S*α*and Sμ-S*ε* DNA in mouse *Rad52^+/+^, Rad52^−/−^* and *Aicda^−/−^* naïve B cells stimulated with nil, LPS only, LPS plus IL-4, LPS plus TGF-*β* and RA, CD154 only, CD154 plus IL-4, or CD154 plus TGF-*β* and RA, as well as Sμ-S*γ*3 in *Rad52^+/+^, Rad52^−/−^* and *Aicda^−/−^*B cells stimulated with LPS only, were analyzed 96 h post stimulation by nested long-range PCR using forward Iμ and reverse Cδ, S*γ*1, S*γ*3, S*α*or S*ε* primers, respectively, followed by Southern-blotting using specific Sμ, σδ, S*γ*1, S*γ*3, S*α*or S*ε* probe, as indicated. Data are one representative of 3 independent experiments yielding comparable results. **b,** IgD titers in culture (96 h) fluid of *Rad52^+/+^* and *Rad52^−/−^*B cells stimulated with LPS plus IL-4, as measured by dot-blotting (two-fold serial diluted culture fluid) using a rat anti-mouse IgD mAb. Data are one representative of 5 independent experiments yielding comparable results.

### Rad52 is required to mount a specific IgD antibody response

We determined the role of Rad52 in supporting a specific IgD antibody response by immunizing *Rad52^−/−^* and *Rad52^+/+^* mice with OVA (20 μg in alum, i.p., 3 times). *Rad52^−/−^* mice showed no Sµ–σδ recombination in spleen, mesenteric lymph nodes (MLNs) or Peyer’s patch B cells (Fig. 6a). The lack of CSR to IgD was specific, as B cells in such mice showed Sµ–S*γ*1 and Sµ– S*α* DNA recombinations as B cells in *Rad52^+/+^* mice, which also underwent Sµ–σδ recombination. In *Rad52^−/−^* mice, Sµ–S*γ*1 and Sµ–S*α* DNA junctions showed fewer and shorter microhomologies than in *Rad52^+/+^* mice (Fig. 6b, Extended Data Figs. 2,3), a reflection of involvement of Rad52 in CSR to isotypes other than IgD^20^. *Rad52^−/−^* mice showed significantly decreased total and/or OVA-specific IgD in circulating blood, bronchoalveolar lavage (BALF), feces (free or bound to fecal bacteria), and IgD-producing cells in MLNs and lamina propria, as compared to their *Rad52^+/+^* counterparts (Fig. 6c-h). This contrasted with the normal or elevated total and OVA-specific IgM, IgG1 and IgA levels in the same *Rad52^−/−^* mice, as predicted based on our previous findings^27^. Thus, Rad52 is required to mount an efficient antigen-specific class-switched IgD response.

**Fig. 6.**
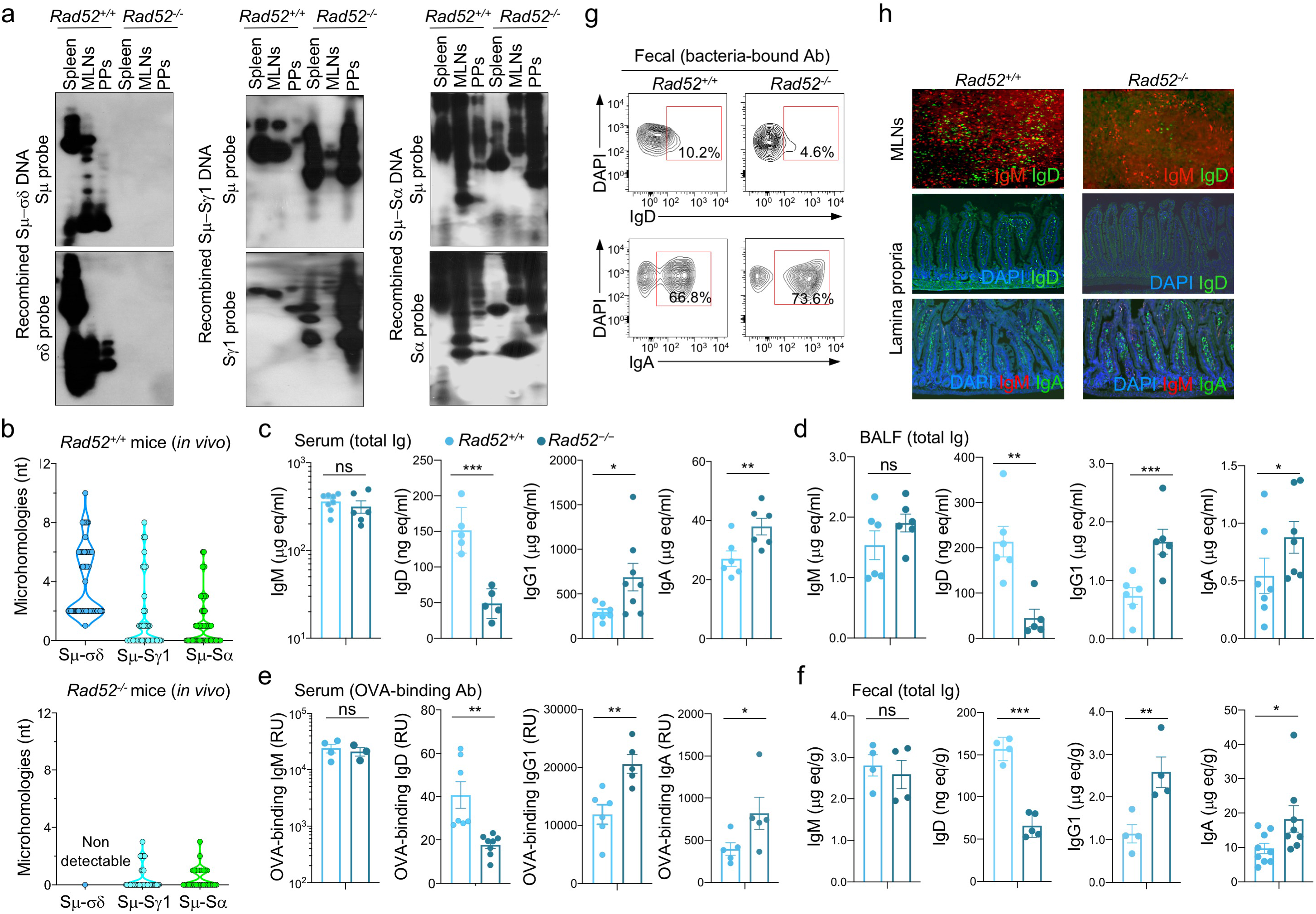
*Rad52* deletion ablates *in vivo* Sμ-σδ DNA recombination and reduces IgD production. *Rad52*^+/+^ and *Rad52^−/−^* mice were immunized (i.p.) with OVA. **a,** Recombined Sμ-σδ, Sμ-S*γ*1, and Sμ-S*α* DNA in spleen, mesenteric lymph nodes (MLN) and Peyer’s patches B cells, as analyzed by nested long-range PCR using forward Iμ and reverse Cδ, S*γ*1 or S*α*primers, followed by Southern-blotting using specific Sμ, σδ, S*γ*1 or S*α* probe, as indicated. Data are one representative of 3 independent experiments yielding comparable results. **b,** Sμ-σδ, Sμ-S*γ*1 and Sμ-S*α* junctional DNAs were amplified by nested PCR and sequenced using by MiSeq. The length and numbers of nucleotide overlaps (microhomologies) in Sμ-σδ, Sμ-S*γ*1 and Sμ-S*α* junctional DNAs are shown by violin plots. Each symbol represents a unique sequence (*n* = 45 per group). **c-f,** Titers of total IgD in serum, BALF and feces, as analyzed by dot-blotting using rat anti-mouse IgD mAb. Titers of total IgM, IgD, IgG1 and IgA as well as OVA-binding IgM, IgD, IgG1 and IgA as analyzed by specific ELISAs. Each dot represents data from one individual mouse (*n* = 5-8 per group, pooled from two experiments). \**p <* 0.05, \*\**p <* 0.01, \*\*\**p <* 0.001, ns: not significant (unpaired *t* test). **g,** Bacteria-bound IgD and IgA in feces as analyzed by flow cytometry. **h,** IgM, IgD and IgA positive cells in mesenteric lymph nodes (MLNs) and lamina propria as visualized by fluorescence microscopy. Data in **g,h** are representative of 3 independent experiments.

### Rad52 is modulated and phosphorylated by IgD CSR-inducing stimuli, and it is recruited to Sμ and σδ

We analyzed *Rad52, Ku70, Ku86* and *Aicda* transcripts as well as respective Rad52, Ku70, Ku86 and AID proteins, including phosphorylated Rad52 (p-Rad52 has been shown to display enhanced ssDNA annealing activity^28^), in B cells induced to undergo CSR to IgD. Mouse B cells stimulated by LPS plus IL-4 and human B cells stimulated by CD154 plus IL-4 and IL-21 increased *Ku70/Ku86* and Ku70/Ku86 expression at 24-48 h concomitant with significantly greater expression of *Aicda* and AID, which was nearly undetectable at time 0, while somewhat downregulating *Rad52* and Rad52. Rad52 protein, however, was progressively phosphorylated within the same time range (Fig. 7a-c). Further supporting its role in CSR to IgD, Rad52 was recruited to Sμ, σδ (and S*γ*1) in B cells stimulated by LPS plus IL-4, which induced Sμ-σδ (and Sμ-S*γ*1) DNA recombination, but not by stimuli that did not induce Sµ–σδ recombination, i.e., LPS alone or LPS plus TGF-*β* and RA, as shown by chromatin immunoprecipitation (ChIP) using an anti-Rad52 Ab – the specificity of the ChIP Rad52 recruitment assay being emphasized by the lack of chromatin immunoprecipitation in *Rad52^−/−^* B cells (Fig. 7d,e). Recruitment of Rad52 but not Ku70/Ku86 to σδ in CSR to IgD, as induced by LPS plus IL-4, contrasted with that of Ku70/Ku86 to S*γ*3 and S*α* regions as induced in CSR to IgG3 and IgA (Fig. 7f), a possible reflection of the competition of these HR and NHEJ elements for binding to S region DSB ends^20^. Notably, LPS plus IL-4 induced recruitment of Rad52 but not Ku70/Ku86 to σδ, while inducing mostly Ku70/Ku86 recruitment to C*γ*1, consistent with the efficient LPS pus IL-4 induction of CSR to IgG1, mediated mainly by NHEJ^20^. Thus, Rad52 expression and, importantly, Rad52 phosphorylation are modulated by IgD CSR-inducing stimuli.

**Fig. 7.**
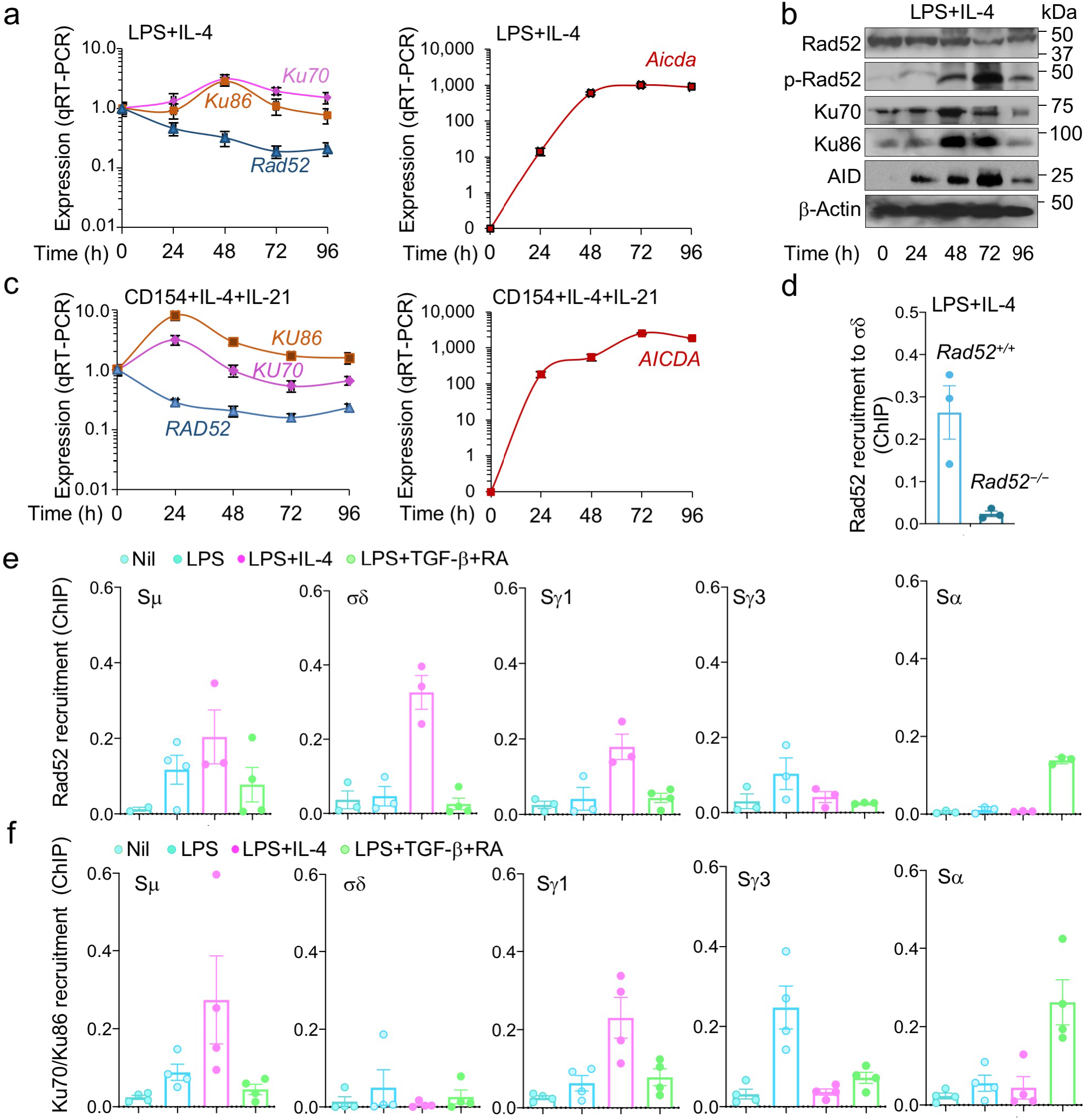
Rad52 is phosphorylated and recruited to Sμ and σδ in B cells induced to undergo IgD CSR. **a,** C57BL/6 mouse naïve B cells were stimulated with LPS plus IL-4 and cultured for 0, 24, 48, 72 and 96 h. *Rad52*, *Ku70*, *Ku86* and *Aicda* transcripts were analyzed by real-time qRT–PCR, normalized to *β-Actin* expression and depicted as relative to the expression in unstimulated B cells (set as 1.0). Data are mean ± SEM of 3 independent experiments. **b,** Expression of Rad52, phosphorylated Rad52 (p-Rad52), AID, Ku70, Ku86, and β-Actin proteins in mouse B cells stimulated with LPS plus IL-4 (as in **a**), as analyzed by specific immunoblotting. Data are one representative of 3 independent experiments yielding comparable results. **c,** Human peripheral blood naive B cells were stimulated with CD154 plus IL-4 and IL-21 and cultured for 0, 24, 48, 72 and 96 h. *RAD52*, *KU70*, *KU86* and *AICDA* transcripts were analyzed by real-time qRT-PCR, normalized to *β-ACTIN* expression and depicted as relative to the expression in unstimulated B cells (set as 1.0). Data are mean ± SEM of 3 independent experiments. **d,** Recruitment of Rad52 to σδ region DNA, as analyzed by ChIP-qPCR assays in mouse *Rad52^+/+^* and *Rad52^−/−^* B cells stimulated with LPS plus IL-4 and cultured for 72 h. Data are expressed as percent of pre-IP input for each sample (mean ± SEM). **e,f,** C57BL/6 mouse B cells were stimulated with nil, LPS, LPS plus IL-4 or LPS plus TGF-*β* and RA and cultured for 72 h. Recruitment of Rad52 (**e**) and Ku70/Ku86 (**f**) to Sμ, σδ, S*γ*1, S*γ*3 and S*α* region DNA, as analyzed by ChIP-qPCR assays. Data are mean ± SEM of 3 independent experiments.

### Stimuli that induce Sμ-σδ DNA recombination downregulate ZFP318/Zfp318 and lead to IgD secretion

Next, we addressed the expression of mIgD and sIgD and its regulation by stimuli inducing CSR to IgD. Resting B cells expressed mIgD and mIgM, but little or no sIgD or sIgM, reflecting high levels of *V_H_DJ_H_-Cδm* and *V_H_DJ_H_-Cμm* trancripts and low levels of *V_H_DJ_H_-Cδs* and *V_H_DJ_H_-Cμs* transcripts (Fig. 8a). Induction of CSR to IgD (by LPS or CD154 plus IL-4) resulted in loss of virtually all mIgD, emergence of *V_H_DJ_H_-Cδ*s transcripts together with *V_H_DJ_H_-Cμs* transcripts and significant IgD secretion (Fig. 8a-b). By contrast, application of IgD CSR non-inducing stimuli (LPS plus TGF-*β* and RA) to similar naive IgM^+^IgD^+^B cells resulted in partial loss of mIgD, no change in *V_H_DJ_H_-Cδm* transcripts and marginal IgD secretion (Fig. 8a-b). The changes in *V_H_DJ_H_-Cδm, V_H_DJ_H_-Cδs* transcripts, mIgD and sIgD brough about by IgD CSR-inducing stimuli paralleled the downregulation of *Zfp318* transcripts and Zfp318 protein – Zfp318 represses the TTS that mediates alternative transcriptional *V_H_DJ_H_-Cμ/V_H_DJ_H_-Cδ* termination, thereby allowing for long-range transcription throughout *V_H_DJ_H_-Cµ-s-m-Cδ-s-m* DNA (Fig. 8c-e). Zfp318 downregulation was specific to IgD CSR, as it did not occur in response to IgA CSR-inducing stimuli (LPS plus TGF-*β* and RA). *ZFP318* downregulation concomitant with decreased mIgD expression and increased IgD secretion was reproduced in human B cells submitted to IgD CSR-inducing stimuli (CpG plus IL-2 and IL-21) but not IgD CSR non-inducing stimuli (CpG plus IL-4 and IL-21) (Fig. 8,f). Similarly, *ZFP318* transcripts and ZFP318 protein were downregulated in human B cells undergoing IgD CSR *in vivo*, as in tonsils (Fig. 8,g). Zfp318 downregulation was independent and likely preceding expression of AID or Rad52, as revealed by virtual absence of *Zfp318* transcripts in LPS plus IL-4-induced *Aicda^−/−^* B cells, *Rad52^−/−^* B cells and *Rad52^+/+^* B cells, all of which lost mIgD expression as compared to similar B cells stimulated by IgA CSR-inducing stimuli (LPS plus TGF-*β* and RA) (Fig. 8h-j). Thus, the stimuli that specifically induce CSR to IgD downregulate ZFP318/Zfp318 independently of AID or Rad52 expression and prior to Sμ-σδ DNA recombination.

**Fig. 8.**
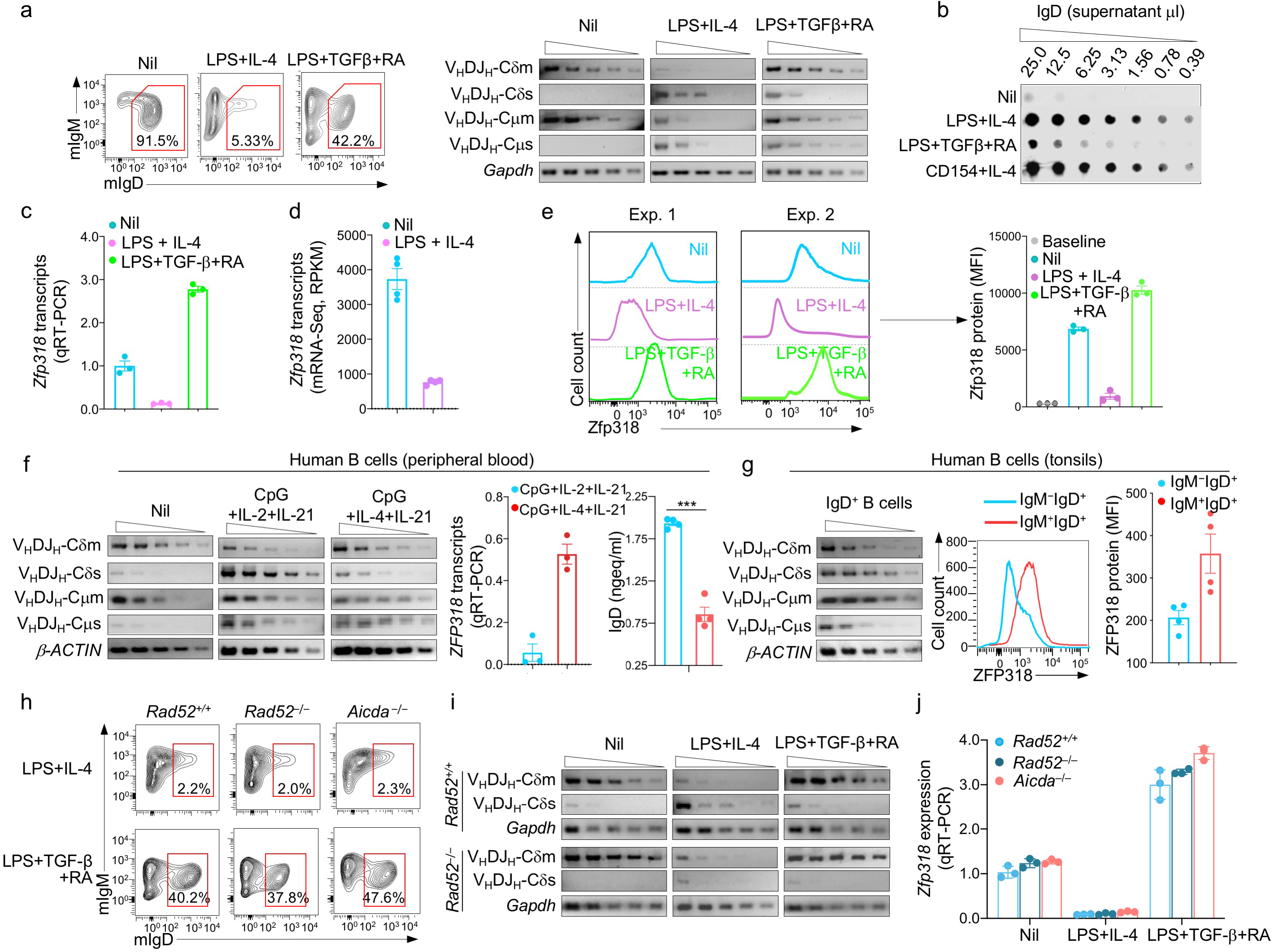
Stimuli inducing Sμ-σδ DNA recombination downregulate ZFP318/Zfp318 in human and mouse B cells. **a**, C57BL/6 mouse naïve B cells were stimulated with nil, LPS plus IL-4 or LPS plus TGF-*β* and RA. Surface expression of IgM and IgD were analyzed 96 h post stimulation by flow cytometry. Expression of *V_H_DJ_H_-Cδ−m*, *V_H_DJ_H_-Cδ−s*, *V_H_DJ_H_-Cμ−m* and *V_H_DJ_H_-Cμ−s* transcripts were analyzed 72 h post-stimulation by semi-quantitative RT-PCR using serial two-fold dilution of cDNA templates. Data are representative of 3 independent experiments. **b,** IgD in supernatant from cultures (96 h) of C57BL/6 mouse naïve B cell stimulated with nil, LPS plus IL-4, LPS plus TGF-*β* and RA, or CD154 plus IL-4, as analyzed by dot-blotting using rat anti-mouse IgD mAb. Data are representative of 5 independent experiments. **c,** Expression of *Zfp318* transcripts in mouse naïve B cells stimulated with nil, LPS plus IL-4, or LPS plus TGF-*β* and RA, as analyzed 72 h post-stimulation by qRT-PCR and normalized to *β-Actin* expression and depicted relative to the average expression in unstimulated B cells (set as 1). Data are mean ± SEM of 3 independent experiments. **d,** Expression of *Zfp318* transcripts in mouse naïve B cells stimulated with LPS plus IL-4, as analyzed 96 h post-stimulation by mRNA-Seq. Data are mean ± SEM of 4 independent experiments. **e,** Zfp318 protein level in mouse naïve B cells stimulated with nil, LPS plus IL-4, or LPS plus TGF-*β* and RA, as analyzed 96 h post-stimulation by intracellular staining with rabbit anti-Zfp318 Ab in flow cytometry. Bars on the right panel represent the MFI (mean ± SEM) from 3 independent experiments. **f,** Human blood naïve B cells stimulated with nil, CpG plus IL-2 and IL-21 or CpG plus IL-4 and IL-21. Human *V_H_DJ_H_-Cδ−m*, *V_H_DJ_H_-Cδ−s, V_H_DJ_H_-Cμ−m* and *V_H_DJ_H_-Cμ−s* transcript levels were measured 72 h post-stimulation by semi-quantitative RT-PCR with serial two-fold dilution of cDNA templates – data are representative of 3 independent experiments (left panels). Expression of *ZFP318* transcripts as analyzed 72 h post-stimulation by qRT-PCR and normalized to *HPRT* expression (**f** middle panel) – data are mean ± SEM of 3 independent experiments. Secreted IgD in supernatants of the human B cell cultures, as analyzed 120 h post stimulation by specific ELISA (**f** right panel) – data are mean ± SEM of 4 independent experiments. **g,** Expression of *V_H_DJ_H_-Cδ−m, V_H_DJ_H_-Cδ−s, V_H_DJ_H_-Cμ−m* and *V_H_DJ_H_-Cμ−s* transcripts in human total IgD^+^ tonsil B cells, as analyzed by semi-quantitative RT-PCR involving serial two-fold dilution of cDNA templates (left panel) – data are representative of 3 independent experiments. Expression of ZFP318 protein in human tonsil IgM^+^IgD^+^B cells and IgM^−^IgD^+^B cells, as analyzed by intracellular staining with anti-Zfp318 Ab in flow cytometry (middle panel) – data are representative of 4 independent experiments (mean ± SEM, right panel). **h,** Surface expression of IgM and IgD in mouse naïve *Rad52^+/+^, Rad52^−/−^* and *Aicda^−/−^*B cells stimulated with LPS plus IL-4, or LPS plus TGF-*β* and RA, as analyzed 96 h post-stimulation by flow cytometry. Data are representative of 3 independent experiments. **i,** Expression of *V_H_DJ_H_-Cδ−m* and *V_H_DJ_H_-Cδ−s* transcripts in mouse naïve *Rad52^+/+^* and *Rad52^−/−^*B cells stimulated with nil, LPS plus IL-4 or LPS plus TGF-*β* and RA, as analyzed 72 h post-stimulation by semi-quantitative RT-PCR using serial two-fold dilution of cDNA templates. Data are representative of 3 independent experiments. **j,** Rad52 or AID deficiency does not alter *Zfp318* expression. Expression of *Zfp318* transcripts in mouse naïve *Rad52^+/+^, Rad52^−/−^* and *Aicda^−/−^*B cells stimulated with nil, LPS plus IL-4, or LPS plus TGF-*β* and RA, as analyzed 72 h post-stimulation by qRT-PCR and normalized to *β-Actin* expression, as depicted relative to expression in unstimulated B cells (set as 1). Data are mean ± SEM of 3 independent experiments.

### RAD52 knockdown reduces Sμ-σδ DNA recombination and IgD secretion in human B cells

The high frequency of microhomologies in Sμ-σδ junctions of human tonsil B cells *in vivo* and human B cells induced to undergo CSR to IgD *in vitro* (Figs. 3a,4,6b, Extended Data Figs. 1-5) suggested to us that RAD52 also mediates Sμ-σδ DNA recombination in human B cells. We purified naïve IgM^+^IgD^+^B cells from peripheral blood of 3 healthy subjects and knocked down using *RAD52*-specific siRNAs *RAD52* transcripts and RAD52 protein by up to 75% and 95%, respectively. In these B cells, Sμ-σδ recombination, as induced by CpG plus IL-2 and IL-21, was virtually abolished, while *AICDA* or AID expression and Sμ-S*γ*1 recombination were not altered (Fig. 9a-c). The reduced Sμ-σδ DNA recombination in RAD52 knockdown human B cells was associated with decreased expression of *V_H_DJ_H_-Cδs* transcripts, without significant alteration of *V_H_DJ_H_-Cδm* transcripts (Fig. 9d). The critical role of RAD52 in human CSR to IgD was emphasized by RAD52 recruitment to Sμ and σδregions in human naïve B cells induced to undergo CSR to IgD (by CpG plus IL-2 and IL-21) *in vitro*, human tonsil (IgD^+^) B cells undergoing CSR to IgD *in vivo*, but not in unstimulated naïve IgD^+^IgM^+^ B cells (Fig. 9e). Thus, Rad52 critically mediates CSR to IgD through Sµ–σδ recombination in human B cells.

**Fig. 9.**
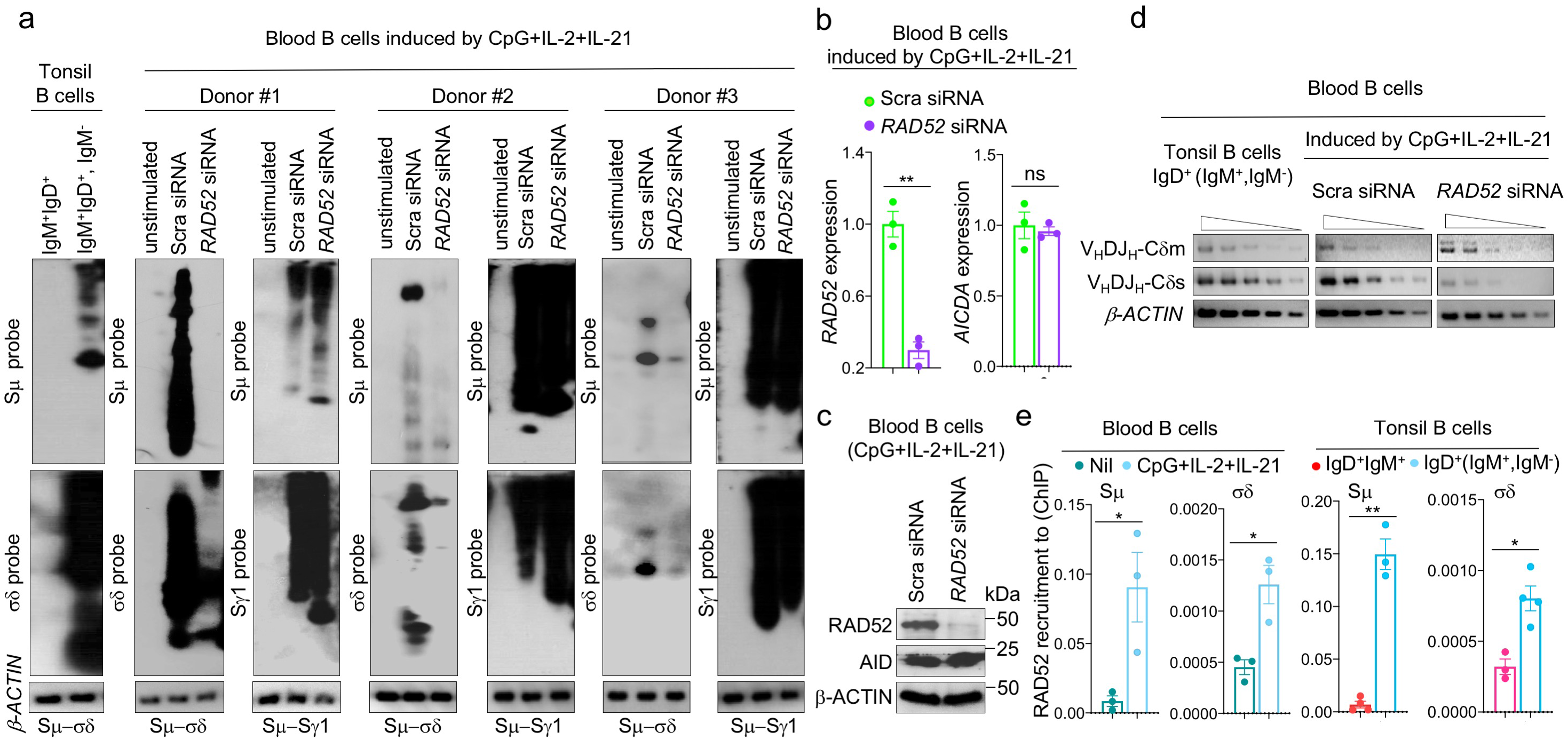
RAD52 is required for Sμ-σδ DNA recombination in human B cells. **a,** Human blood naïve IgM^+^IgD^+^B cells were transfected with specific *RAD52* siRNA or scrambled (Scra) siRNA and stimulated by CpG plus IL-2 and IL-21. Recombined Sμ-σδ and Sμ-S*γ*1 DNA in the transfected B cells 120 h after siRNA transfection, as well as Sμ-σδ DNA in tonsil IgM*^−^*IgD^+^ and blood naive IgM^+^IgD^+^B cells were analyzed by nested long-range PCR using forward Iμ and reverse Cδ or S*γ*1 primers followed by Southern-blotting using indicated specific probes. Data are from 3 independent experiments. **b**, Expression of *RAD52* and *AICDA* transcripts was analyzed 48 h after siRNA transfection by qRT-PCR and normalized to *HPRT* expression. Data are mean ± SEM of 3 independent experiments. **c**, Expression of RAD52 and AID proteins were analyzed 72 h after siRNA transfection by specific immunoblotting. Data are representative of 3 independent experiments. **d**, Expression of *V_H_DJ_H_-Cδm* and *V_H_DJ_H_-Cδs* transcripts as analyzed by semi-quantitative RT-PCR with of serial two-fold dilution of cDNA templates. Data are representative of 3 independent experiments. **e,** RAD52 is recruited to σδ region DNA in human B cells undergoing CSR to IgD. Recruitment of RAD52 to Sμ and σδ region DNA in human blood B cells stimulated for 120 h with CpG plus IL-2 and IL-21, as analyzed by specific ChIP-qPCR. Data are mean ± SEM of 3 or 4 independent experiments.

### Sμ-σδ DNA recombination leads to IgD plasma cell differentiation

To determine whether the substantial IgD production we observed only upon induction of CSR to IgD (Figs. 5b,6c-h,8b,f) would be associated with plasma cell differentiation, we analyzed human IgM*^+^*IgD^+^B cells induced to undergo CSR to IgD by CpG plus IL-2 and IL-21. More than 13% of these B cells became mIgM^−^intracellular IgD^+^ compared to about half of their counterparts stimulated by CpG plus IL-4 and IL-21 and not switching to IgD (Fig. 10a). More than 90% of the IgM^−^IgD^+^B cells emerging from CpG plus IL-2 and IL-21 simulation expressed BLIMP-1 and almost 60% were CD27^+^CD38^+^ versus about 10% of the IgM^−^IgD^+^B cells from CpG plus IL-4 and IL-21 expressing BLIMP-1 and less than 12% being CD27^+^CD38^+^. Among mouse IgM*^+^*IgD^+^B cells induced to undergo CSR to IgD by LPS plus IL-4, about 25% expressed intracellular IgD. All these B cells also expressed Blimp-1 and 70% or more acquired CD138 (Fig. 10b). By contrast, among IgM*^+^*IgD^+^B cells induced to undergo CSR to IgA by LPS plus TGF-*β* and RA, about 50% expressed intracellular IgD but virtually none expressed Blimp-1 or acquired surface CD138. The relevance of IgD CSR to sustained IgD secretion was suggested by analysis of 3 human myelomas, two IgD and one IgA. Both IgD myelomas displayed Sµ–σδ DNA, but not Sµ–S*α* DNA recombination (Fig. 10c). Conversely, the IgA myeloma showed Sµ–S*α*, but not Sµ–σδ DNA recombination. Thus, IgD^+^B cells emerging by CSR would are prone to differentiate into IgD-secreting plasmablasts/plasma cells for sustained IgD secretion. And such IgD^+^B cells may function as precursors of neoplastic IgD^+^ transformants.

**Fig. 10.**
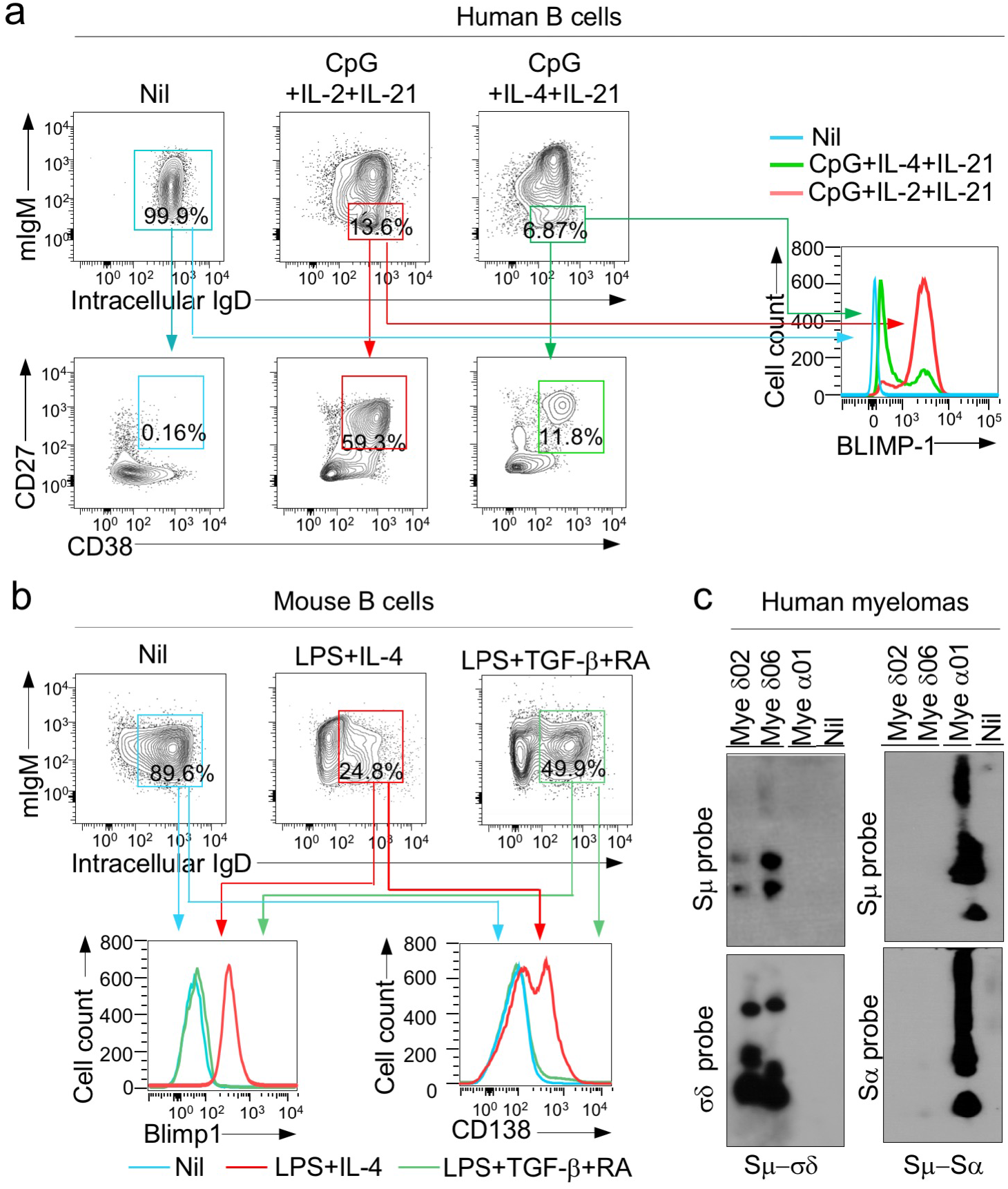
B cells undergoing CSR to IgD differentiate to IgD-producing plasmablasts/plasma cells. **a,** Human B cells stimulated with CpG plus IL-2 and IL-21, which induce IgD CSR, or CpG plus IL-4 and IL-21, which do not induce IgD CSR. Proportions of CD138^+^IgD*^+^*IgM*^−^* plasmablasts/plasma cells among intracellular sIgM*^−^* IgD*^+^*B cells and Blimp1 expression in intracellular IgD*^+^* sIgM*^−^* cells, as analyzed 120 h post stimulation by flow cytometry. **b,** mouse *Rad52^+/+^* and *Rad52^−/−^*B cells stimulated with LPS plus IL-4, which induce IgD CSR. Proportions of sCD138^+^ plasmablasts/plasma cells among intracellular IgD*^+^* sIgM*^−^* cells and Blimp1 expression in intracellular IgD*^+^* sIgM*^−^* cells, as analyzed 96 h post stimulation by flow cytometry. Data in **a** and **b** are representative of 3 independent experiments. **c,** Recombined Sμ-σδ and Sμ-S*α* DNA in two IgD^+^ myelomas and one IgA^+^ myeloma as analyzed by nested long-range PCR followed by Southern-blotting using indicated probes.

### B cell Rad52 phosphorylation, increased CSR to IgD and IgD autoantibodies in systemic autoimmunity

Serum IgD have been suggested to increase in patients with inflammatory autoimmune diseases, such as systemic lupus erythematosus (SLE)^29^ and rheumatoid arthritis^30^, and in hereditary autoinflammatory syndromes, most notably the hyper-IgD syndrome (HIDS)^31–34^. While in healthy humans, many B cells make IgD that react with components of the self^35^, we found patients with systemic lupus to display significantly higher levels of circulating IgD, including IgD specific for nuclear antigens, than their healthy subject controls (Fig. 11a,d). This possibly reflected the higher level of B cell Rad52 and/or p-Rad52 expression in such lupus patients (Fig. 11k). Similarly, we found lupus-prone MRL/*Fas^lpr/lpr^* mice to show far higher levels of IgD than their wildtype C57BL/6 counterparts in serum, feces and BALF as well as increased IgD-coated bacteria in feces, a reflection of high levels of *V_H_DJ_H_-Cδs* transcripts in bone marrow, spleen, MLNs and Peyer’s patches B cells as well as increased numbers of IgD^+^ B cells in lamina propria, MLNs and Peyer’s patches (Fig. 11a-f). In MRL/*Fas^lpr/lpr^* mice, the elevated IgD levels reflected increased IgD-producing cells and increased Sμ-σδ DNA recombination in bone marrow, spleen, MLNs and Peyer’s patches (Fig. 11g). Increased CSR to IgD in MRL/*Fas^lpr/lpr^* was associated with high levels of p-Rad52 expression, greater frequency and length of microhomologies in Sµ–σδ as compared to Sµ-S*γ*1 and Sµ-S*α* junctional sequences, as well as with a high frequency of somatic point-mutations in areas abetting Sμ-Sδ DNA junctions (Fig. 11h-k and Extended Data Fig. 6). Thus, high levels of B cells expressing p-Rad52 are associated with high levels of IgD and IgD autoantibodies to nuclear antigens in lupus patients and in lupus-prone MRL/*Fas^lpr/lpr^* mice. In these mice, Sμ-σδ DNA recombination events involving high frequency of junctional microhomologies occur in B cells of different body districts, giving rise to high levels of IgD autoantibodies locally and systemically.

**Fig. 11.**
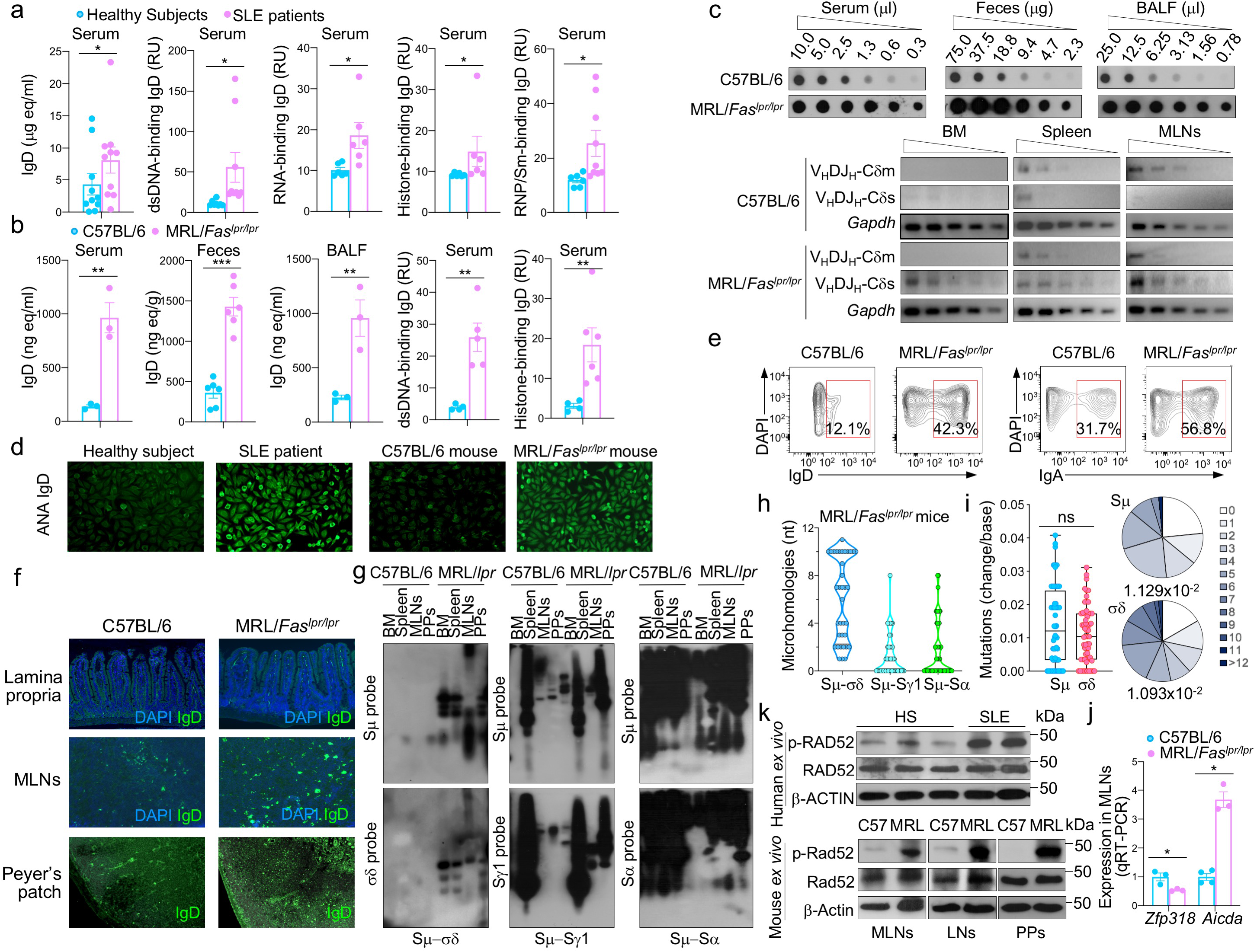
p-Rad52 expression, CSR to IgD and anti-nuclear antigen IgD autoantibodies in lupus mice and patients. **a,** Concentrations of total, and dsDNA-, RNA-, histone- or RNP/Sm-binding IgD in healthy human subjects and SLE patients, as analyzed by specific ELISAs. Data are mean ± SEM of 6 to 10 healthy subjects or SLE patients. **b,** Concentrations of total IgD in serum, feces and BALF as analyzed by dot-blotting, and concentrations of dsDNA- or histone-binding IgD autoantibodies in serum of C57BL/6 and MRL/*Fas^lpr/lpr^* mice as analyzed by specific ELISAs. Data are mean ± SEM of 3 to 9 mice. **c,** IgD concentrations in the serum, BALF and feces of C57BL/6 and MRL/*Fas^lpr/lpr^* mice, as analyzed by dot-blotting. Shown are dot-blots from one C57BL/6 and one MRL/*Fas^lpr/lpr^* mouse, representative of 3 to 9 C57BL/6 and MRL/*Fas^lpr/lpr^* mice. Expression of *V_H_DJ_H_-Cδm, V_H_DJ_H_-Cδs, V_H_DJ_H_-Cμm* and *V_H_DJ_H_-Cμs* transcripts in bone marrow (BM), spleen and mesenteric lymph nodes (MLNs) as analyzed semi-quantitative RT-PCR in serial two-fold dilutions of cDNA templates. Shown are RT-PCR data from one C57BL/6 and one MRL/*Fas^lpr/lpr^* mouse, representative of 3 C57BL/6 and 3 MRL/*Fas^lpr/lpr^* mice. **d,** ANAs as visualized by indirect immunofluorescence on HEp-2 cells that were incubated with sera from a human healthy subject and a SLE patient or C57BL/6 and MRL/*Fas^lp/lpr^* mice and revealed using FITC-labeled rat mAb to mouse IgD. **e,** Bacteria-bound IgD and IgA in feces of C57BL/6 and MRL/*Fas^lpr/lpr^* mice, as analyzed by flow cytometry. **f,** IgD^+^B cells in lamina propria, MLNs and Peyer’s patches of C57BL/6 and MRL/*Fas^lpr/lpr^* mice, as visualized by fluorescent microscopy. **g,** Recombined Sμ-σδ, Sμ-S*γ*1, and Sμ-S*α* DNA in bone marrow, spleen, MLNs and Peyer’s patches B cells from C57BL/6 and MRL/*Fas^lpr/lpr^* mice were analyzed by nested long-range PCR using forward Iμ and reverse Cδ, S*γ*1 or S*α* primers, followed by Southern-blotting using indicated probes. Data are representative of 3 independent experiments. **h,** Sμ-σδ, Sμ-S*γ*1 and Sμ-S*α* junctional DNAs in non-immunized MRL/*Fas^lpr/lpr^* mice, as amplified by nested PCR and sequenced by MiSeq. The length and numbers of nucleotide overlaps (microhomologies) in Sμ-σδ, Sμ-S*γ*1 and Sμ-S*α* junctional DNAs are shown by violin plots. Each dot represents a unique sequence (*n* = 45 per group). **i**, Somatic point-mutations in Sμ and σδ regions abetting recombined Sμ*−σδ*DNA junctions in IgD class-switched spleen B cells from 3 MRL/*Fas^lpr/lpr^* mice. Mutations were identified in a 48 to 506 nt stretch of Sμ or σδ regions in unique Sμ-σδ DNA recombination sequences. Each dot represents an individual sequence. In pie charts, the size of slices denotes the proportion of sequences with the same number of mutations and the grey hue denotes the number of point-mutations per sequence; below the pie charts is the overall mutation frequency (change/base). \*\**p <* 0.05, \*\**p <* 0.01, \*\*\**p <* 0.001, ns: not significant (unpaired *t* test). **j,** Expression of *Zfp318* and *Aicda* transcripts in MLNs from non-immunized C57BL/6 and MRL/*Fas^lpr/lpr^* mice as analyzed by specific qRT-PCR. Data are mean ± SEM of 3 C57BL/6 and 3 MRL/*Fas^lpr/lpr^* mice. **k,** Expression of phosphorylated Rad52 (p-Rad52), Rad52 and β-Actin proteins in peripheral blood B cells from healthy human subjects and SLE patients as well as B cells from non-immunized C57BL/6 and MRL/*Fas^lpr/lpr^* mice, as analyzed by specific Western blotting using rabbit anti-p-Rad52 Ab (Y408472, Applied Biological Materials Inc.) or anti-β-Actin mAb (2F1-1, BioLegend). P-Rd52 (Y104) Ab detected endogenous levels of Rad52 protein only when phosphorylated at tyrosine 104.

## Discussion

The mechanism of CSR to IgG, IgA and IgE are quite well understood, as mediated by Ku70/Ku86-dependent NHEJ, although occurrence of a “residual” IgM to IgG CSR in B cells lacking Ku70/Ku86 expression has suggested the existence of a Ku70/Ku86-independent A-EJ synaptic mechanism^18–20^. Mice lacking 53BP1, in which NHEJ-dependent CSR to IgG, IgA and IgE was significantly decreased – 52BP1 protects resection of DSB ends, thereby skewing the synaptic process toward NHEJ – showed increased CSR to IgD and increased circulating IgD levels, suggesting that the short-range Sµ–σδ CSR was mediated by a 53BP1-independent synaptic process involving resected DSB ends and entailing a high frequency of Sµ–σδ junctional microhomologies^22, 23^. This together with our previous demonstration that Rad52 plays a central role in synapsing intra-Sμ region resected DSB ends as well as *c-Myc/IgH* locus translocations also involving resected DSB ends, both processes entailing significant junctional microhomologies, prompted us to hypothesize that Rad52 mediates the A-EJ process that synapses Sµ and σδ DSB with complementary overhangs in CSR to IgD^20^. Here, we demonstrated that Rad52 mediates CSR to IgD in mouse and human B cells (Extended Data Fig. 7), thereby unveiling a previously unknown, critical and dedicated role of this HR factor in mammalian DNA repair.

We have provided here unequivocal evidence that Rad52 is critical for CSR to IgD *in vitro* and *in vivo,* in mouse and human B cells. In mouse *Rad52^−/−^* B cells, Sμ-σδ DNA recombination was ablated and IgD secretion greatly reduced. Similarly, in *RAD52* knockdown B cells from healthy subjects, Sμ-σδ DNA recombination was virtually abrogated and IgD secretion greatly decreased – as expected^1, 5^, Sμ-σδ DNA recombination could not occur in the absence of AID, which introduces DSBs in σδ as it does in Sμ, S*γ*, S*α* or S*ε*. In mouse *Rad52^−/−^* B cells and human *RAD52* knockdown B cells, decreased post-recombination *V_H_DJ_H_-Cδs* transcripts resulted in reduced IgD secretion, which occurred in presence of unaltered transmembrane *V_H_DJ_H_-Cδm* transcript levels, at least within the first 72 hours from CSR induction. Interestingly, the stimuli that selectively induced CSR to IgD modulated the overall levels of *Ku70/KU70*, *Ku86/KU86* and *Rad52/RAD52* transcripts while significantly upregulating *Aicda/AICDA* in mouse and human B cells. This was concomitant with induction of AID and moderate decrease in Rad52 protein, which, in fact, was increasingly phosphorylated at Tyr104. Rad52 Tyr104 phosphorylation has been shown to boost Rad52-mediated DNA single-strand annealing and is possibly effected by c-ABL kinase^28^. Rad52 involvement in CSR to IgD was further emphasized by recruitment of this protein to σδ region (in addition and necessarily to Sμ) *in vivo* in human tonsil IgD^+^ B cells, as well as *in vitro*, in mouse and human naïve B cells induced to undergo CSR to IgD, but no or only marginally in similar B cells undergoing CSR to IgG3 or IgA. Instead, these B cells recruited Ku70/Ku86 to S*γ*3 and S*α*, consistent with the major contribution of NHEJ to CSR to IgG3 and IgA.

Rad52 is a member of the eponymous epistasis group for DSB repair that shows strong evolutionary conservation^17, 36^. In *Saccharomyces cerevisiae,* Rad52 is a key element of the HR pathway, and its deletion or mutation impairs DNA DSB repair^37, 38^. Indeed, yeast Rad52 is a recombination mediator and a facilitator of annealing of complementary DNA single-strands^39, 40^. It functions as a cofactor of Rad51, which forms nucleoprotein filaments with single-strand DNA and promotes strand pairing, by overcoming the inhibitory effect of replication protein A (RPA)^41^. By contrast, Rad52 mutation or even deletion results in no obvious abnormalities in viability or functions in mammalian cells. As we have shown, *Rad52^−/−^* mice displayed no significant alteration of immune system elements, including B cells^20^, possibly owing to the presence of mammalian gene paralogues, such as *BRCA2* and *RAD51*, which by encoding functions related to Rad52, can compensate for the absence of this factor^42^. Human BRCA2 functions as a recombination mediator by facilitating RAD51 nucleoprotein filament formation^40, 43–45^. Nevertheless, human BRCA2 cannot facilitate annealing of RPA-coated DNA, a function that Rad52 carries out efficiently in the absence of BRCA2^46^. This together with Rad52 involvement in DSB repair at stalled or collapsed replication forks points at a unique role of Rad52 in catalyzing single-strand annealing in homology-directed DNA repair in human cells^47–49^.

Our identification of Rad52 as essential in IgD CSR Sμ-σδ synapses provides, to the best of our knowledge, the first demonstration of a critical and dedicated role of this factor in mammalian DNA repair. The short-range Rad52-mediated Sμ-σδ recombination of resected DSB ends adds to the other Rad52-mediated short-range DSB recombination we recently uncovered: intra-Sμ region DSB recombination^20^. This, like Sμ*−σδ* synapsing, engages resected DSB ends and yields significant junctional nucleotide microhomologies^20^. In this function, as in CSR to IgD, Rad52 is not fungible in mouse or human B cells. Our identification of Rad52 as the critical element in Sμ-σδ synapsis also sheds light on the mechanistic nature of the CSR A-EJ DSB repair pathway (originally referred to as A-NHEJ^19^). As per our current findings, the CSR A-EJ pathway uses HR Rad52 to synapse upstream and downstream DSB overhangs by a MMEJ process but does not require a homologous template as a guide, as the HR pathway does. The DNA polymerase *θ* has been suggested to contribute to A-EJ^21^. Our previous findings, however, did not support a role of this polymerase in Rad52-mediated intra-Sμ DSB recombination or Sμ*−*S*γ*1, Sμ*−*S*γ*3, Sμ*−*S*γ*2a/S*γ*2c and Sμ*−*S*α* recombination (CSR to IgG1, IgG3, IgG2a/IgG2c and IgA)^20^. Finally, while (MMEJ) A-EJ functions as a back-up pathway in cells defective in NHEJ or HR, it also synapses DSB ends in cells that are competent for both NHEJ and HR^50^, as exemplified by microhomologies in S-S junctions in a proportion of B cells that class-switched to IgG, IgA and IgE, as well as the disappearance of such microhomologies upon Rad52 ablation^20^.

As we showed here, Rad52 works in concert with Zfp318 to modulate IgD expression through an interplay of alternative RNA splicing and DNA recombination, the latter after AID intervention. Zfp318 represses the TTS intercalated between the Cμ and Cδ exons within the *Ighμ/Ighδ* loci transcriptional complex unit^10, 11^, thereby allowing for transcription of long primary *V_H_DJ_H_-Cµ-s-m-Cδ-s-m* RNA. Zfp318, however, would also simultaneously allow for the continuous expression of primary *Iµ-Cµ-s-m-Cδ-s-m* RNA transcripts. In fact, albeit possibly more abundant, hence their predominant detection in our specific PCR assays, *Iµ-Cδ* transcripts – in secretory or membrane form – are identical in sequence to their post-recombination *Iµ-Cδ* counterparts (Fig. 1a,b). During B cell development, Zfp318 expression closely parallels mIgD expression^10, 11^. Indeed, consistent with its repression of the TTS intercalated between the *Cμ* and *Cδ* exons complex, the Zfp318 protein is expressed during the transition from immature IgM^+^IgD^−^ to mature IgM^+^IgD^hi^ B cell^10, 11^. As we showed here, naïve mature B cells which express high levels of mIgD also express high levels of *Zfp318* transcripts and Zfp318 protein. In these B cells, stimuli that induced Sμ-σδ DNA recombination, yielding primary *V_H_DJ_H_-Cδ-s-m* RNA transcripts, also induced profound downregulation of *Zfp318* transcripts and Zfp318 protein, suggesting that relieving Zfp318-mediated TTS repression is a prerequisite for Sμ-σδ DNA recombination to unfold. Conversely, as we also showed here, naïve mature IgM^+^IgD^+^ B cells submitted to stimuli that induced CSR to isotypes other than IgD, such as IgA (by LPS plus IL-4 and RA), further upregulated *Zfp318* transcripts and Zfp318 protein, concomitant with no Sμ-σδ recombination, thereby allowing for massive expression of mIgD rather than sIgD.

The role of Zfp318 as gene transcription regulator is highly specific for IgD, as genome-wide transcriptome analysis of B cell Zfp318-deficient (*Vav-Cre* dependent deletion) mice identified *Sva* as the only other gene altered in expression^11^ – interestingly, *Sva* is also involved in alternative splicing, albeit outside the *IgH* locus^51^. Zfp318 would be under the control of 5’AMP-activated protein kinase (Ampk). This is phosphorylated by Lbk1^52^, whose signaling triggers the B cell GC reaction. Indeed, Lbk1’s failure to activate Ampk or Ampk loss specifically muted Zfp318 expression and IgD transcription^52^. In contrast, activation of Ampk by phenformin impaired GC formation^52^, likely by heightening Zfp318 expression, possibly in addition to other mechanisms. This would result in increased expression of primary *V_H_DJ_H_-Cµ-s-m-Cδ-s-m* RNA transcripts but not Sμ-σδ DNA recombination, suggesting that CSR to IgD is one of the multiple and complex events inherent to GC formation. This is triggered by naturally occurring generally microbial stimuli, as in tonsil GCs and GCs or other secondary lymphoid formations in aerodigestive mucosae^1, 5, 6^. Consistent with the contention that Ampk mediates the regulation of Zfp318 as well as the contrasting impact of IgD CSR-inducing (LPS plus IL-4) and non-inducing stimuli (LPS plus TGF-*β* and RA) on expression of Zfp318, stimulation of both human and mouse cells by LPS has been shown to result in dephosphorylation/inactivation of Ampk, while similar cell stimulation by TGF-*β* resulted in rapid phosphorylation/activation of this protein kinase^53^.

In our experiments, stimuli that induced CSR to IgD (e.g., LPS plus IL-4 in mouse B cells, and CD154 plus IL-2 and IL-21 in human B cells) also downregulated Zfp318 expression which, in turn, reduced *V_H_DJ_H_-Cδm* transcript level and mIgD, while greatly increasing *V_H_DJ_H_-Cδs* transcripts and sIgD. This argues for CSR to IgD to be critical for significant IgD secretion. Indeed, stimuli that induced Sμ-σδ recombination and IgD secretion also induced plasmablast/plasma cell differentiation, as shown by Blimp-1 and C38^+^CD27^+^ expression in human B cells, and Blimp-1 and CD138^+^ in mouse B cells. A similar outcome was not produced by stimuli that did not induce Sμ-σδ recombination and IgD secretion in mouse or human B cells. Thus, while alternative splicing of long primary *V_H_DJ_H_-Cµ-s-m-Cδ-s-m* RNA transcripts in B cells that have not undergo CSR would make some contribution to the overall level of IgD production *in vivo*, CSR to IgD is likely required for substantial and sustained IgD production, as secreted by plasmablasts/plasma cells or by neoplastic transformants, such as IgD myeloma cells. The limited IgD amounts detected in supernatants of mouse or human B cells primed by stimuli that induced high levels of mIgD but not Sµ–σδ synaptic recombination would result from translation of alternative spliced long primary *V_H_DJ_H_-Cµ-s-m-Cδ-s-m* RNA transcripts as well as some “shedding” of mIgD.

Bacteria and viruses have been suggested to play an important role in driving CSR to IgD, generally through stimulation of TLRs in gut and respiratory lymphoid tissues, and mesenteric lymph nodes, possibly leading to emergence of plasmablasts and plasma cells secreting IgD^1, 3–5, 51, 54–56^. Circulating IgD are increased in patients with frequent respiratory infections or chronic lung inflammation suggesting a protective role for this Ig isotype^1, 3, 5, 56^. Our findings support the notion that both T-dependent (CD154) and T-independent (TLR ligands) stimuli induce Sμ-σδ DNA recombination, in combination with various cytokines^1, 4–6^. Interestingly, although we previously showed that BCR-signaling synergizes with TLR-signaling for induction of AID and CSR to IgG and IgA^25^, BCR signaling did not synergize with TLR7 or TLR9 signaling to induce CSR to IgD, as shown by the lack of Sμ-σδ DNA recombination in B cells stimulated by CpG or R848 plus IL-4 and anti-Igδ Ab. In the *in vivo* T-dependent antibody response to OVA, ablation of CSR to IgD (*Rad52^−/−^* mice) resulted in reduced levels of total and specific IgD in circulating blood and BALF, decreased total and/or bacteria-bound IgD in feces as well as decreased numbers of IgD^+^ B cells lamina propria and MLNs, a privileged site of IgD CSR^12^. As predicted by our previous findings^20^, the overall decreased IgD levels in *Rad52^−/−^* mice were associated with increased IgG1 and IgA as well as greatly decreased frequency and lengths of microhomologies in Sμ-S*γ*1 and Sμ-S*α* junctions. This reflected the lack of Rad52 contribution to the synaptic process underpinning such junctions as well as the lack of Rad52 competition with Ku70/Ku86^20^, which resulted solely in Ku70/Ku86-mediated NHEJ, a process that limits microhomologies to 0-3 nt^16^.

Information on the contribution of IgD to autoimmunity is scant and contradictory. Self-antigen-binding and mostly polyreactive IgD occur in healthy subjects, much like IgM or even IgG and IgA do^35, 57–60^. High levels of IgD have been reported in rheumatoid arthritis patients and thought to possibly be implicated in the pathogenesis of the disease^30^. mIgD expression, however, has been speculated to exert an inhibitory effect on B cell autoreactivity, as suggested by elevated autoantibody production, increased deposition of immune complexes in kidneys and severe nephritis in lupus-prone C56BL/6*lpr* mice with deletion of the Igδ locus^61, 62^. Our findings showed total and self-reactive IgD (dsDNA, histone, RNP/Sm or RNA and ANAs) to be elevated in the circulation of lupus patients and lupus-prone MRL/*Fas^lpr/lpr^* mice. The latter displayed higher levels of IgD in serum, BALF and feces, than their wildtype C57BL/6 counterparts. Such high IgD levels reflected CSR recombinations that included Sμ-σδ junctions with extensive microhomologies and high frequency of somatic mutations in the DNA areas abetting Sμ-σδ junctions. Such IgD CSR occurred in different districts, such as bone marrow, spleen, MLNs and Peyer’s patches, and were reflected in the IgD^+^B cells in those districts. This together with the B cell high levels of p-Rad52 and the low levels Zfp318 indicated that in murine and likely human lupus, IgD autoantibodies stem from extensive B cell Sμ-σδ recombination rather than alternative splicing of primary *V_H_DJ_H_-Cµ-s-m-Cδ-s-m* RNA transcripts. Our findings do not suggest a “protective” role of IgD in autoimmunity^61, 62^, while supporting a role of CSR to IgD in systemic lupus autoantibody responses.

Collectively, our data outline a critical and dedicated role of Rad52 in mediating the synapsis of Sμ with σδ DSB resected ends. They also provide the first demonstration of Rad52 as a critical element in the poorly understood contribution of A-EJ to the resolution of DSBs in nonmalignant cells. In malignant B cells, Rad52 is involved in DNA recombination events that give rise to DNA deletions and translocations. As we previously showed, Rad52 ablation reduced the frequency of *c-Myc/IgH* translocations in mouse *p53*^−/−^ B cells by more than 70%, with the residual translocations containing limited microhomologies^20^. Whether Rad52 intervention extends to other modalities of A-EJ in neoplastic and non-neoplastic lymphoid mammalian cells remains to be determined. The importance of this newly unveiled and essential function of Rad52 is further emphasized by our demonstration that this highly conserved HR element is critical for CSR to IgD in both mouse and human B cells. This together with the further reduction of the physiologically moderate microhomologies in Sµ–S*γ*1, Sµ–S*γ*3, Sµ–S*α* and Sµ– S*ε* junctions in *Rad52^−/−^* B cells (current data and refs.^18–20^) solidifies the role of Rad52 as critical mediator of the A-EJ backup pathway underpinning the residual CSR to IgG, IgA and IgE in the absence of Ku70/Ku86 proteins^20^. Our findings also showed how stimuli that induce Sμ-σδ recombination coordinate Rad52 function, as enabled by phosphorylation, with downregulation of Zfp318, unique repressor of the TTS intercalated between the Cμ and Cδ loci, whose activity allows transcription throughout *V_H_DJ_H_-Sμ-Cµ-s-m-σδ-Cδ-s-m* and *Iµ-Cµ-s-m-σδ-Cδ-s-m*. Further, they indicate that CSR to IgD is required for sustained IgD secretion and possibly a prerequisite for IgD plasma cell differentiation. They also add new and significant information to a potential role of CSR to IgD, as promoted by Rad52 phosphorylation, in systemic autoimmunity. Finally, they provide important new molecular information to approach the virtually unexplored mechanistic underpinning of hyper-IgD syndrome, a relatively rare but a severe autoinflammatory disease associated with mevalonate kinase deficiency (due to *MVK* recessive mutations) and exorbitant levels of IgD^31, 32, 63^.

## Acknowledgements

We thank Dr. Patrick M. Sung for reviewing this manuscript. We also would like to thank Amanda Fisher, Dr. Justin B. Moroney, Dr. Helia N. Sanchez and Dr. Huoqun Gan for their help in some experiments. This work was supported by NIH grants R01 AI 079705, T32 AI138944, R01 AI 105813 and the Lupus Research Alliance Target Identification in Lupus Grant 641363 to P.C.

## Author contributions

Y. Xu and H. Zhou performed experiments; G. Post provided myeloma samples; H. Zan designed and performed experiments, analyzed data, supervised the work and wrote the manuscript; P. Casali planned the study, designed the experiments, analyzed the data, supervised the work and wrote the manuscript.

## Methods

### Mice

*Rad52^−/−^* mice were generated by replacing exon 3 of the *Rad52* gene with positive selection marker neomycin, as driven by the phosphoglycerate kinase (PGK) promoter, and an upstream mouse sequence functioning as a transcription terminator (Dr. Albert Pastink, Leiden University, Leiden, The Netherlands)^36^. *Rad52^−/−^* mice were backcrossed to C57/BL6 mice for more than six generations. No full length or truncated Rad52 protein was produced from the disrupted allele^36^. *Rad52^−/−^* mice were viable and fertile, and showed no gross abnormalities. *Aicda^−/−^* mice (C57BL/6 background)^64^ were obtained from Dr. Tasuku Honjo (Kyoto University, Kyoto, Japan). C57BL/6 and MRL/*Fas^lpr/lpr^* mice were purchased from Jackson Laboratory (Bar Harbor, Maine). All mice were housed in pathogen-free conditions. Both male and female mice aged 8-12 weeks were used for the experiments. The Institutional Animal Care and Use Committees (IACUC) of the University of Texas Health Science Center at San Antonio approved all animal protocols.

### Mouse B cells and CSR induction *in vitro*

Naïve IgM^+^IgD^+^B cells were isolated from spleens of 8–12-week-old C57BL/6, *Rad52^−/−^* or *Aicda^−/−^* mice as described^25^. B cells were resuspended in RPMI 1640 medium with 10% FBS (FBS-RPMI), 50 mM β-mercaptoethanol and 1x antibiotic-antimycotic mixture (15240-062; Invitrogen) and stimulated with LPS (4 μg/ml) from *Escherichia coli* (055:B5; Sigma-Aldrich), CD154 (1 U/ml; obtained from membrane fragments of baculovirus-infected Sf21 insect cells^25^), CpG ODN 1826 (1.0 μM; Eurofins Genomics) or R848 (1.0 μM; Medkoo) plus nil, IL-4 (5.0 ng/ml; R&D Systems) and/or TGF-β (2.0 ng/ml; R&D Systems) and retinoic acid (RA, 10 nM) or anti-BCR Ab (anti-δ mAb-dextran, 30 ng/ml; Fina Biosolutions). Mouse B cells were cultured in FBS-RPMI at 37°C in 48-well plates for 24, 48, 72 and 96 h.

### Human B cells and CSR induction *in vitro*

Naïve IgM^+^IgD^+^B cells were purified by negative selection using the EasySep^TM^ human naive B cell enrichment kit (19254; StemCell Technologies) from healthy subject PBMCs, following manufacturer’s instructions. Tonsillar IgD^+^ B cells were isolated from human tonsil cells by positive selection using biotin-anti-human IgD mAb (clone IA6-2; 348212, Biolegend) and MagniSort™ Streptavidin Positive Selection Beads (MSPB-6003-74, Thermo Fisher Scientific). Naïve B cells were stimulated with CD154 (10 U/ml) or CpG ODN 2395 (1.0 μM; Eurofins Genomics) plus nil, IL-2 (20 ng/ml; BioLegend), IL-4 (20 ng/ml; R&D Systems), IL-15 and/or IL-21 (50 ng/ml; R&D Systems). Human B cells were cultured in FBS-RPMI at 37°C in 48-well plates for 24, 48, 72, 96 and 120 h.

### Flow cytometry

For surface staining, mononuclear cells were reacted with VF-anti-CD19 mAb (75-0193-0100, Tonbo), PE-anti-IgM mAb (clone RMM1, 406507, BioLegend), and FITC anti-mouse IgD mAb (clone 11-26c.2a, 405704, BioLegend) and 7-AAD. For intracellular staining, cells were stained with anti-CD19 mAb (Clone 1D3; Tonbo) and fixable viability dye eFluor^®^ 450 (FVD 450, eBiosciences) followed by incubation with the BD Cytofix/Cytoperm buffer at 4°C for 20 min. After washing twice with the BD Perm/Wash buffer, cells were resuspended in HBSS with 1% BSA and stored overnight at 4°C. Cells were then stained with anti-Zfp318 Ab (AAS23325C, Antibody Verify; labeled with FITC using iLink™ Antibody Labeling Kits, ABP Biosciences). FACS analysis was performed on single cell suspensions. In all flow cytometry experiments, cells were appropriately gated on forward and side scattering to exclude dead cells and debris. Cell analyses were performed using a LSR-II flow cytometer (BD Biosciences), and data were analyzed using FlowJo software (TreeStar). All experiments were performed in triplicates.

### Fluorescence microscopy

Fluorescence microscopy of tissues. To analyze IgM and IgD-producing cells in the lamina propria and PPs, the intestine was folded into a “Swiss-roll”, fixed with PFA (4%), and embedded in paraffin. Ten μm sections were cut and heated at 80 °C to adhere to the slide, washed four times in xylene for 2 min, dehydrated two times with 100% ethanol for 1 min, two times with 95% ethanol for 1 min and washed two times in water for 1 min. Antigens were unmasked using 2 mM EDTA in 100 °C for 40 mins followed by a cooling step at 25 °C on the bench top, 3 times washing with 1x TBS and blocking using 10% BSA for 15 min. Slides were again washed 3 times with 1x TBS and stained with FITC–anti-IgD mAb (clone 11-26c.2a; 405713, BioLegend), PE goat-anti-mouse-IgM mAb (406507, BioLegend) for 2 h in a dark moist chamber. After washing 3 times with Triton X-100 (0.1%) in TBS, slides were air dried, and cover slips were mounted with ProLong^®^ Gold Antifade Reagent with DAPI (Invitrogen). Fluorescence images were captured using a 10x objective lens with a Zeiss Axio Imager Z1 fluorescence microscope. To analyze IgD-producing cells in MLNs, 10 µm MLN sections were prepared by cryostat and loaded onto positively charged slides, fixed in cold acetone and stained with FITC–anti-IgD mAb (405704, BioLegend), or PE goat-anti-mouse-IgM mAb (406507, BioLegend), respectively, for 1 h at 25 °C in a moist chamber. Cover slips were then mounted using ProLong^®^ Gold Antifade Reagent using DAPI (Thermo Fisher), before examination with a fluorescence microscope.

Fluorescence microscopy of B cells. B cells were suspended at 10^5^ cells/100 µl in FCS-RPMI. Pre-labeled slides were then placed into Cytofunnels and ran with 50 µl of FCS-RPMI in order to wet the Cytofunnel paper. Cells were then placed again into the Cytofunnel and spun at 800 RPM for 3 min using a Cytospin™ 4 Cytocentrifuge (Thermo Fisher). For intracellular visualization of IgM, IgD, CD138 and Blimp-1 proteins, cells were fixed with methanol for 15 min and washed 3 times in PBS-Tween 20. Cells were then blocked in 10% BSA for 15 min and stained with 1:20 APC-anti-mouse IgD Ab (clone 11-26c.2a; 405713, BioLegend), FITC-anti-mouse IgM mAb (11-5790-81, Thermo Fisher), overnight in a dark moist chamber.

### Detection of free and bacterial-bound antibodies

Titers of serum, BALF or fecal total IgD, IgM, IgG1 and IgA and OVA-binding IgD, IgM, IgG1 and IgA were measured using specific ELISAs, as we described^20, 25, 65, 66^. Total IgD in *in vitro* culture supernatants of stimulated human and mouse B cells or in serum, BALF or feces were measured by dot blotting with serially two-fold diluted samples.

Bacteria-bound IgD and IgA were detected in feces by flow cytometry, as we described^27^. Feces (10 mg) were suspended in 100 μl 1x PBS (filtered through 0.2 μm filter), homogenized and centrifuged at 400 × g for 5 min to remove large particles. The supernatant was then centrifuged at 8000 g for 10 min to remove non-bound antibodies (in supernatant). The bacterial pellet was suspended in 1 ml of PBS with 1% (w/v) BSA. After fixation with 7.2% formaldehyde for 10 min at room temperature, bacteria were washed with PBS, and stained with FITC– anti-IgD mAb (clone 11-26c.2a; 405713, BioLegend) or FITC-anti-IgA mAb (C10-3, BD Biosciences) on ice for 30 min, washed with PBS, and further resuspended in 1 x PBS containing 0.2 μg ml^-1^ DAPI for flow cytometry analysis. All events that stained with DAPI were considered as bacteria.

### S-S region DNA recombinations and S region somatic mutations

Genomic DNA was prepared from human or mouse B cells using QIAmp DNA Mini Kit (Qiagen), or from paraffin-embedded human IgD or IgA myeloma tissue sections (obtained from the University of Arkansas for Medical Science) using Quick-DNA™ FFPE Kit (Zymo Research). Recombined Sμ–σδ, Sμ–S*γ*1, Sμ–S*α* and Sμ–S*ε* DNA were amplified by two sequential rounds of specific PCR using Phusion™ high-fidelity DNA polymerase (Thermo Scientific™) and nested oligonucleotide primers^67^ (**Supplementary Table 1**). The first and second rounds of PCR were performed at 98 °C for 30 sec, 58 °C for 45 sec, 72 °C for 4 min (30 cycles). Amplified DNA was fractionated through 1.0% agarose, blotted onto Hybond-N^+^ membranes (GE Healthcare) and hybridized to biotin-labeled Sμ and σδ, S*γ*1, S*α*or S*ε* specific probes. Detection was performed using the Chemiluminescent Nucleic Acid Detection Module (Thermo Fisher Scientific) according to the manufacturer’s instructions. For sequence analysis of the recombined DNA, PCR products were purified using a QIAquick PCR purification kit (Qiagen). The amplified library was tagged with barcodes for sample multiplexing, and PCR was enriched and annealed to the required Illumina clustering adapters. High-throughput 300–base pair (bp) paired-end sequencing was performed by the UTHSCSA Genome Sequencing Facility using the Illumina MiSeq platform. S-S junctions and somatic mutations in the S regions were analyzed by sequence alignment as performed by comparing PCR products sequences with germline Sµ and σδ, S*γ*1 or S*α*sequences using National Center for Biotechnology Information BLAST (www.ncbi.nih.gov/BLAST).

### RT-PCR and quantitative RT-PCR (qRT-PCR)

For quantification of mRNA, germline I_H_-C_H_, post-recombination Iμ-C_H_ and mature V_H_DJ_H_-C_H_ transcripts, RNA was extracted from 0.2-5.0 x 10^6^ cells using either Trizol^®^ Reagent (Invitrogen) or RNeasy Plus Mini Kit (Qiagen). Residual DNA was removed from the extracted RNA with gDNA eliminator columns (Qiagen). cDNA was synthesized from total RNA with the SuperScript*™* IV First-Strand Synthesis System (Thermo Fisher) using oligo-dT primer. Transcript expression was measured by qRT-PCR with the appropriate primers (Supplemental Table 1) using a Bio-Rad MyiQ*™* Real-Time PCR Detection System (Bio-Rad Laboratories) to measure SYBR Green (IQ*™* SYBR*^®^* Green Supermix, Bio-Rad Laboratories) incorporation with the following protocol: 95°C for 15 sec, 40 cycles of 94°C for 10 sec, 60°C for 30 sec, 72°C for 30 sec. Data acquisition was performed during 72°C extension step. Melting curve analysis was performed from 72°C-95°C. Mature *V_H_DJ_H_-Cμm*, *V_H_DJ_H_-Cμs*, *V_H_DJ_H_-Cδm* and *V_H_DJ_H_-Cδs* transcripts were analyzed by semi-quantitative PCR using serially two-fold diluted cDNA.

### Western blotting

B cells were lysed in Laemmli buffer. Cell extracts containing equal amounts of protein (50-100 µg) were fractionated through SDS-PAGE (6%). The fractionated proteins were transferred onto polyvinylidene difluoride membranes (Bio-Rad) overnight (30 V/90 mA) at 4 °C. After blocking and overnight incubation at 4 °C with anti-AID antibody (H-80, Santa Cruz), anti-Ku70 antibody (A0883, Abclonal), anti-Ku86 antibody (A5862, Abclonal), anti-Rad52 antibody (H-300, Santa Cruz Biotechnology), anti-phospho-Rad52 antibody (Y408472, Applied Biological Materials Inc.) or anti-β-Actin mAb (2F1-1, BioLegend), the membranes were incubated with horseradish peroxidase (HRP)-conjugated secondary antibodies. After washing with TBS– Tween 20 (0.05%), bound HRP-conjugated antibodies were detected using Western Lightning Plus-ECL reagents (PerkinElmer Life and Analytical Sciences).

### ChIP and qPCR

ChIP assays were performed as previously described^68–70^. Human or mouse B cells (1.0 x 10^7^) were treated with formaldehyde (1% v/v) for 10 min at 25°C to crosslink chromatin, washed once in cold PBS with protease inhibitors (Roche) and resuspended in lysis buffer (20 mM Tris-HCl, 200 mM NaCl, 2 mM EDTA, 0.1% w/v SDS and protease inhibitors, pH 8.0). Chromatin was fragmented by sonication (DNA fragments of about 200 to 1,000 bp in length), pre-cleared with protein A agarose beads (Pierce) and incubated with agarose conjugated anti-Rad52 mAb (clone F-7; sc-365341 AC, Santa Cruz Biotechnology) at 4°C overnight. Immune complexes were washed and eluted (50 mM Tris-HCl, 0.5% SDS, 200 mM NaCl, 100 µg/ml proteinase K, pH 8.0), followed by incubation at 65°C for 4 h. DNA was purified using a QIAquick PCR purification kit (Qiagen). The Sμ or σδ region DNA was amplified from immunoprecipitated chromatin by qPCR using appropriate primers (**Supplemental Table 1**). Data were normalized to input chromatin DNA and depicted as relative abundance of each amplicon.

### *RAD52* knockdown in human B cells

The human RAD52-specific siRNA oligo duplex (TT320001, Locus ID 5893) and non-effective Trilencer-27 Flurescent-labeled transfection control siRNA duplex (SR30002) were obtained from Origene Technologies. The siRNA duplexes were used to transfect purified human naïve B cells using the Human B Cell Nucleofector^TM^ Kit (VPA-1001, LONZA). Transfected B cells were then stimulated with CpG ODN 2395 plus IL-2 and IL-21 for 96 h before genomic DNA extraction for analysis of Sμ-σδand Sμ-S*γ*1 DNA recombination. Expression of *RAD52* and *AICDA* transcripts were analyzed by qRT-PCR using specific primers 24 h after transfection. Expression of RAD52, phosphorylated-RAD52, AID and *β*-ACTIN proteins were analyzed by immune-blotting 24 h after transfection.

### High-throughput mRNA-Seq

RNA was isolated from cells using the Directzol RNA Microprep Kit (Zymogen Research), according to manufacturer’s instructions and as previously described^66^. RNA integrity was verified using an Agilent Bioanalyzer 2100 (Agilent). Next generation RNA-Seq for mRNA and non-coding RNA was performed by the Genome Sequencing Facility at University of Texas Health Science Center San Antonio Greehey Children’s Cancer Research Institute. High-quality RNA was processed using an Illumina TruSeq RNA sample prep kit v2 or TruSeq Small RNA Sample Prep kit following the manufacturer’s instructions (Illumina). Clusters were generated using TruSeq Single-Read Cluster Gen. Kit v3-cBot-HS on an Illumina cBot Cluster Generation Station. After quality control procedures, individual mRNA-Seq or small RNA-Seq libraries were then pooled based on their respective 6-bp index portion of the TruSeq adapters and sequenced at 50 bp/sequence using an Illumina HiSeq 3000 sequencer. Resulting reads were checked by assurance (QA) pipeline and initial genome alignment (Alignment). After the sequencing run, demultiplexing with CASAVA was employed to generate the Fastq file for each sample. All sequencing reads were aligned with their reference genome (UCSC mouse genome build mm9) using TopHat2 default settings, and the Bam files from alignment were processed using HTSeq-count to obtain the counts per gene in all samples. Quality control statistical analysis of outliers, intergroup variability and distribution levels, were performed for statistical validation of the experimental data.

### Statistical analysis

Statistical analysis was performed using Excel (Microsoft) or Prism^®^ GraphPad software. *P*-values were determined by paired and unpaired Student’s *t-*tests; and *P*-values <0.05 were considered significant.

### IRB for use of human tissues and peripheral blood as well as IACUC for use of mice

For the use of DNA procured from formalin fixed paraffin embedded tissues obtained from the University of Arkansas for Medical Science, the study was reviewed by the University of Arkansas for Medical Sciences Institutional Review Board (IRB) which determined that this project is not human subject research as defined in 45 CFR 46.102. Human B cells were purified from PBMCs of healthy subject buffy coats obtained from South Texas Blood and Tissue Center, San Antonio, Texas, under the *Healthy Volunteer Blood Donor Program* and lupus patients B cells were purified PBMCs obtained under the Long School of Medicine IRB HSC 20140234H *Class switching, somatic hypermutation and plasma cell differentiation in B cells.* Mouse and mouse B cell studies were performed under the Long School of Medicine IACUC 20200019AR *Somatic hypermutation, class switch DNA recombination and plasma cell differentiation in antibody and autoantibody responses*.

---

**Extended Data Fig. 1.**
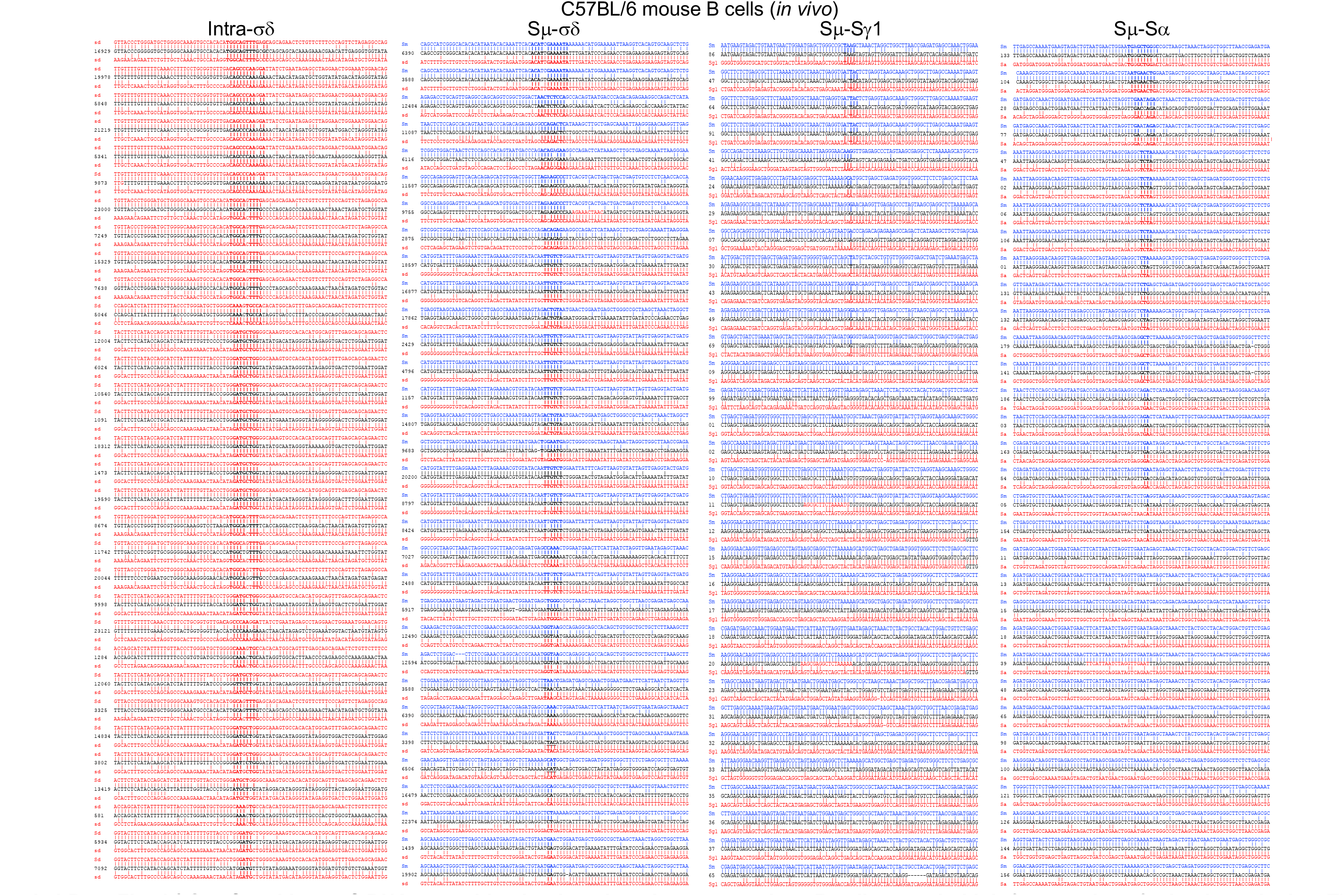
Sμ-σδ and intra-σδ DNA recombination junctions in mouse spleen B cells contain high frequencies of microhomologies. The junctions of intra-σδ, Sμ-σδ, Sμ-Sγ1 and Sμ-Sα recombinant DNAs from spleen B cells of an OVA-immunized C57BL/6 mouse were amplified and sequenced using MiSeq system. Thirty-two representative junction sequences from each group are shown. Each intra-σδ recombinant DNA sequence (middle) is compared with the upstream (above) and the downstream (below) germline σδ sequences. Each Sμ-σδ, Sμ-Sγ1 and Sμ-Sα recombinant DNA sequence (middle) is compared with the germline Sμ (above) and the σδ, Sγ1 or Sα (below) sequence. Microhomologies (bold) were determined by identifying the longest regions at the junctions of perfect uninterrupted donor/acceptor identity or the longest overlap region at the S–S junction with no more than one mismatch on either side of the breakpoint.

**Extended Data Fig. 2.**
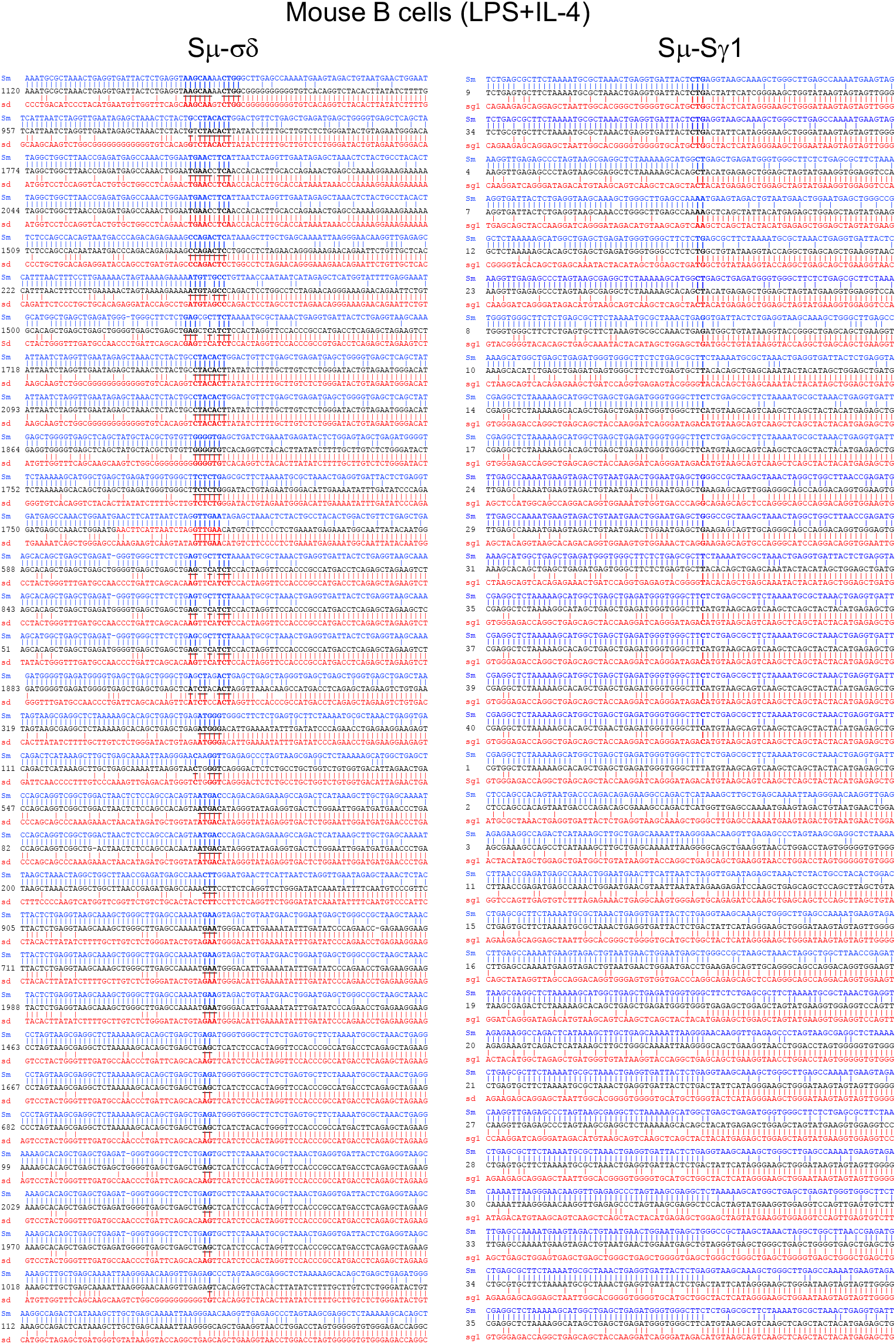
Recombined Sμ−σδ DNA junctions in mouse B cells stimulated *in vitro* to undergo IgD CSR contain high frequencies of microhomologies. C57BL/6 mouse B cells were stimulated with LPS plus IL-4 and cultured for 96 h. The recombined Sμ−σδ and Sμ-Sγ1 DNA junctions were amplified and sequenced using MiSeq system. Thirty-two representative Sμ-σδ and 32 representative Sμ-Sγ1 junction sequences are shown. Each recombinant DNA sequence (middle) is compared with the germline Sμ (above, blue) and the σδ or Sγ1 (below, red) sequence. Microhomologies (bold) were determined by identifying the longest region at the Sμ-σδ or Sμ-Sγ1 junction of perfect uninterrupted donor/acceptor identity or the longest overlap region at the S–S junction with no more than one mismatch on either side of the breakpoint.

**Extended Data Fig. 3.**
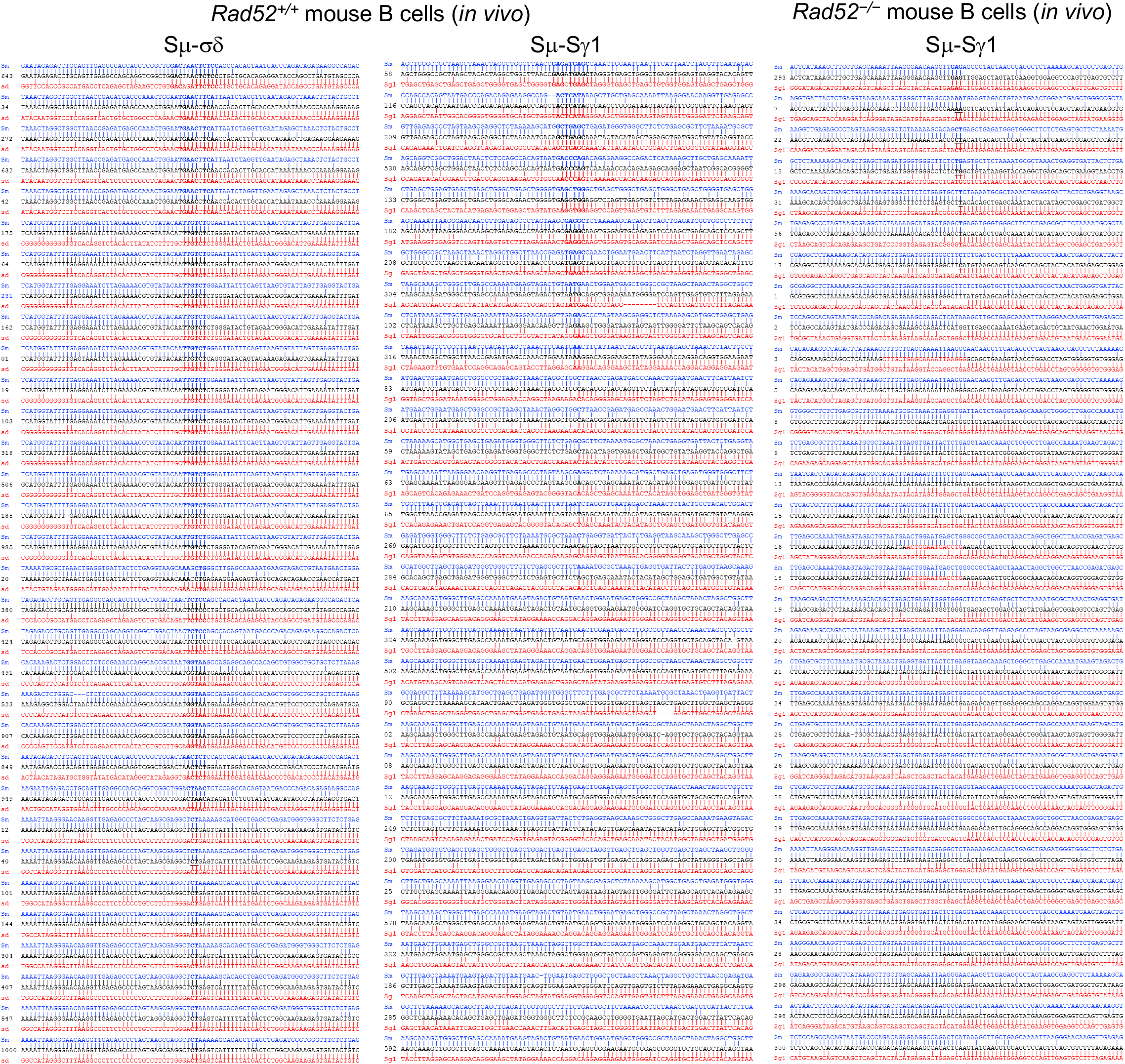
*Rad52–/–* mice display no Sμ−σδ recombination and even lower frequencies of microhomologies in Sμ-Sγ1 junctions. The junctions of recombined Sμ−σδ and Sμ-Sγ1 DNAs from spleen B cells of OVA-immunized *Rad52+/+* and *Rad52–/–* mice were amplified and sequenced using MiSeq system. No Sμ−σδ recombination was detected in B cells from *Rad52–/–* mice. Thirty-two representative Sμ-σδ and 32 representative Sμ-Sγ1 junction sequences are shown. Each recombinant DNA sequence (middle) is compared with germline Sμ (above, blue) and σδ or Sγ1 (below, red) sequences. Microhomologies (bold) were determined by identifying the longest region at the Sμ-σδ or Sμ-Sγ1 junction of perfect uninterrupted donor/acceptor identity or the longest overlap region at the S-S junction with no more than one mismatch on either side of the breakpoint.

**Extended Data Fig. 4.**
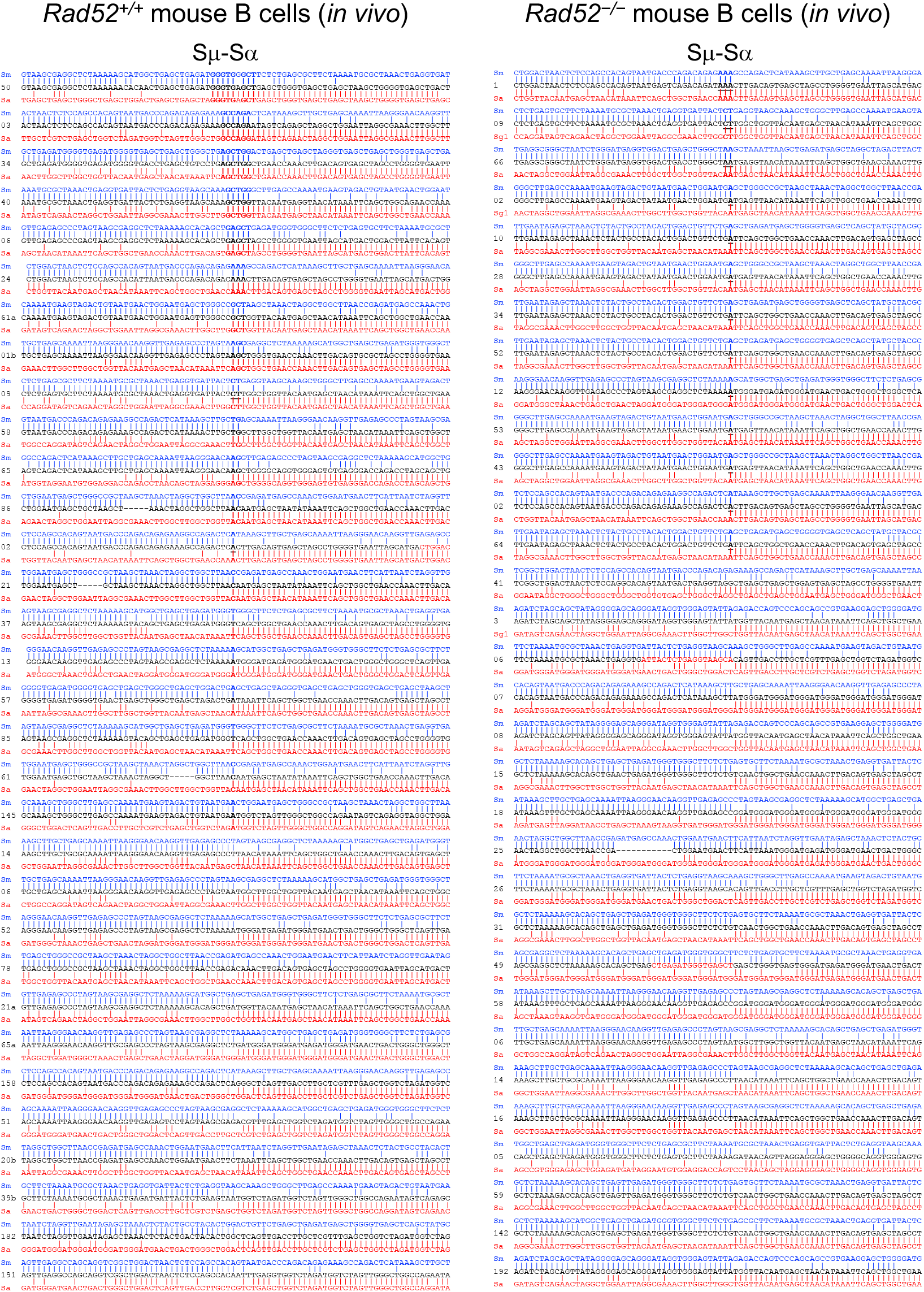
*Rad52–/–* mice display lower frequencies of microhomologies in B cell Sμ-Sα junctions. The junctions of recombined Sμ-Sα DNAs from spleen B cells of OVA-immunized *Rad52+/+* and *Rad52–/–* mice were amplified and sequenced using MiSeq system. Thirty-two representative junction sequences from *Rad52+/+* mice and 32 representative sequences from *Rad52–/–* mice are shown. Each recombinant DNA sequence (middle) is compared with germline Sμ (above, blue) and Sα (below, red) sequences. Microhomologies (bold) were determined by identifying the longest region at the Sμ-Sα junction of perfect uninterrupted donor/acceptor identity or the longest overlap region at the S-S junction with no more than one mismatch on either side of the breakpoint.

**Extended Data Fig. 5.**
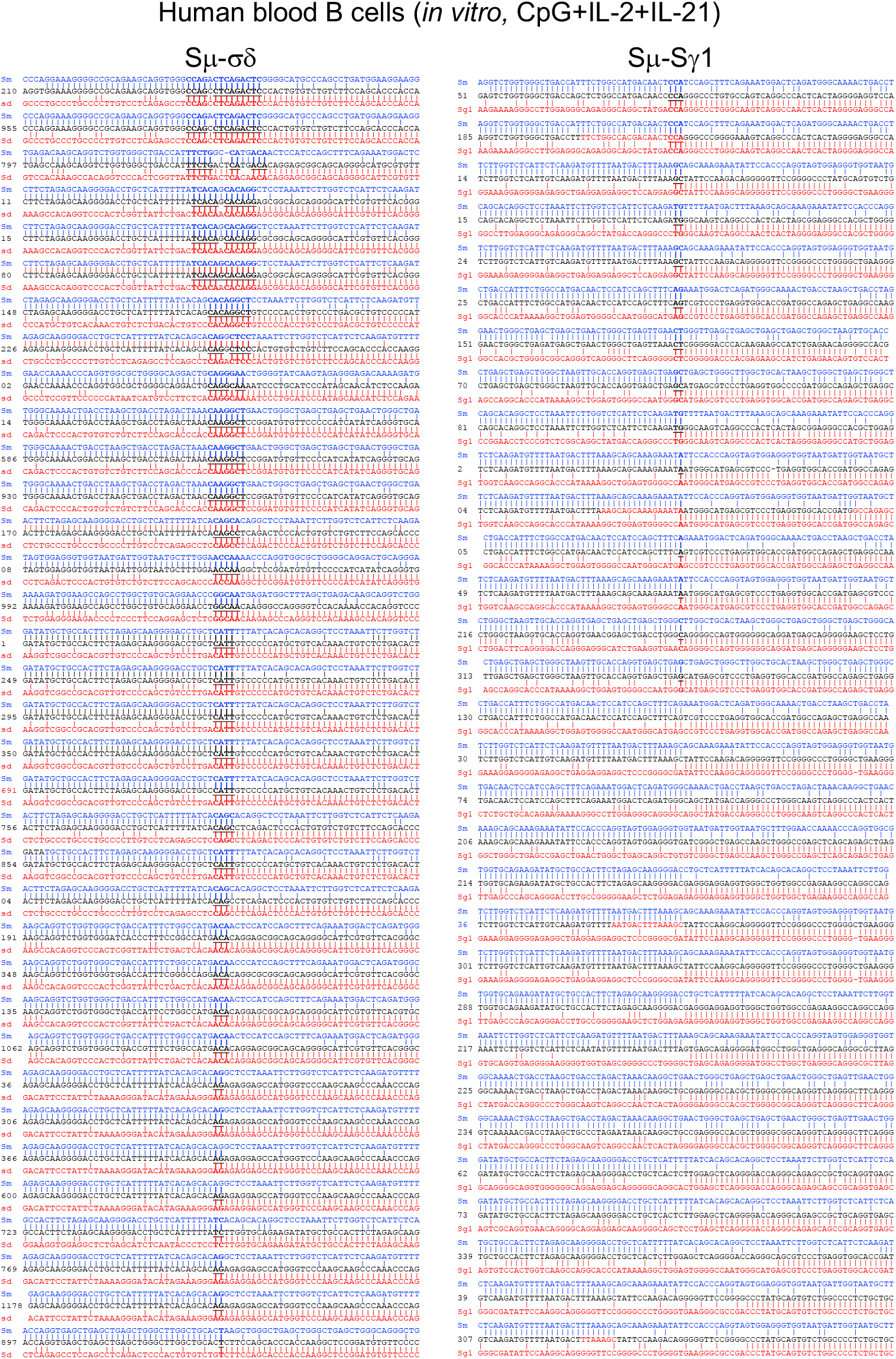
Human B cells stimulated to undergo CSR to IgD and IgG1 *in vitro* display a higher frequency of microhomologies in Sμ-σδ DNA junctions. Human naïve B cells were stimulated with CpG plus IL-2 and IL-21 and cultured for 120 h. The junctions of recombined Sμ-σδ and Sμ-Sγ1 DNAs were amplified and sequenced using MiSeq system. Thirty-two representative sequences from Sμ-σδ and Sμ-Sγ1 junctions are shown. Each recombinant DNA sequence (middle) is compared with germline Sμ (above, blue) and σδ or Sγ1 (below, red) sequences. Microhomologies (bold) were determined by identifying the longest region at the Sμ-σδ or Sμ-Sγ1 junction of perfect uninterrupted donor/acceptor identity or the longest overlap region at the S-S junction with no more than one mismatch on either side of the breakpoint.

**Extended Data Fig. 6.**
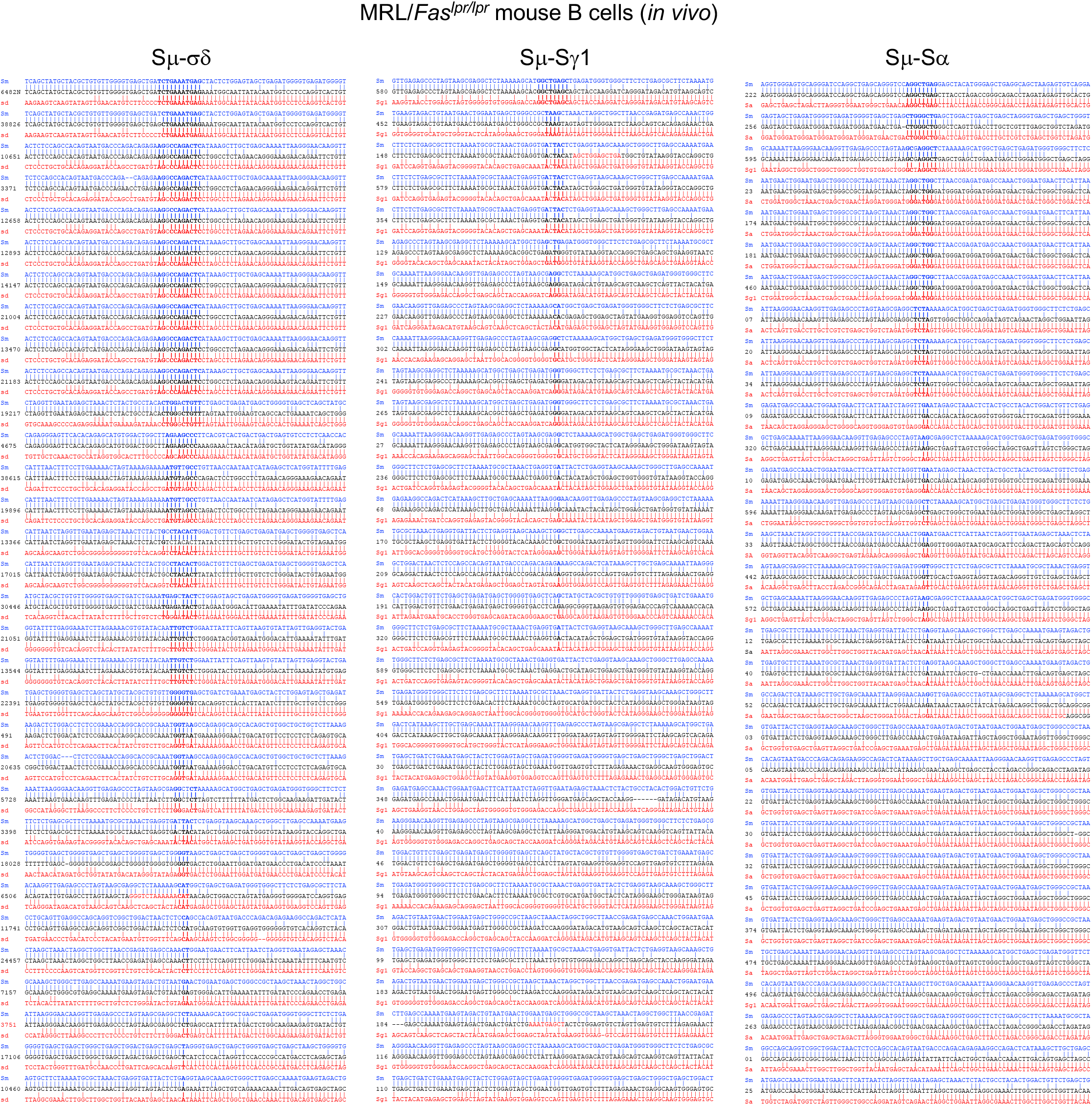
Increased CSR to IgD associates with high frequency of microhomologies in recombined Sμ-σδ, Sμ-Sγ1 and Sμ-Sα DNA junctions in autoimmune mice. The junctions of Sμ-σδ, Sμ- Sγ1 and Sμ-Sα recombinant DNAs from spleen B cells of MRL/*Faslpr/lpr* mice were amplified and sequenced by MiSeq. Thirty-two representative sequences from Sμ-σδ, Sμ-Sγ1 and Sμ-Sα junctions are shown. Each recombinant DNA sequence (middle) is compared with the germline Sμ (above, blue) and σδ, Sγ1 or Sα (below, red) sequences. Microhomologies (bold) were determined by identifying the longest region at the Sμ-σδ, Sμ-Sγ1 or Sμ-Sα junction of perfect uninterrupted donor/acceptor identity or the longest overlap region at the S-S junction with no more than one mismatch on either side of the breakpoint.

**Extended Data Fig. 7.**
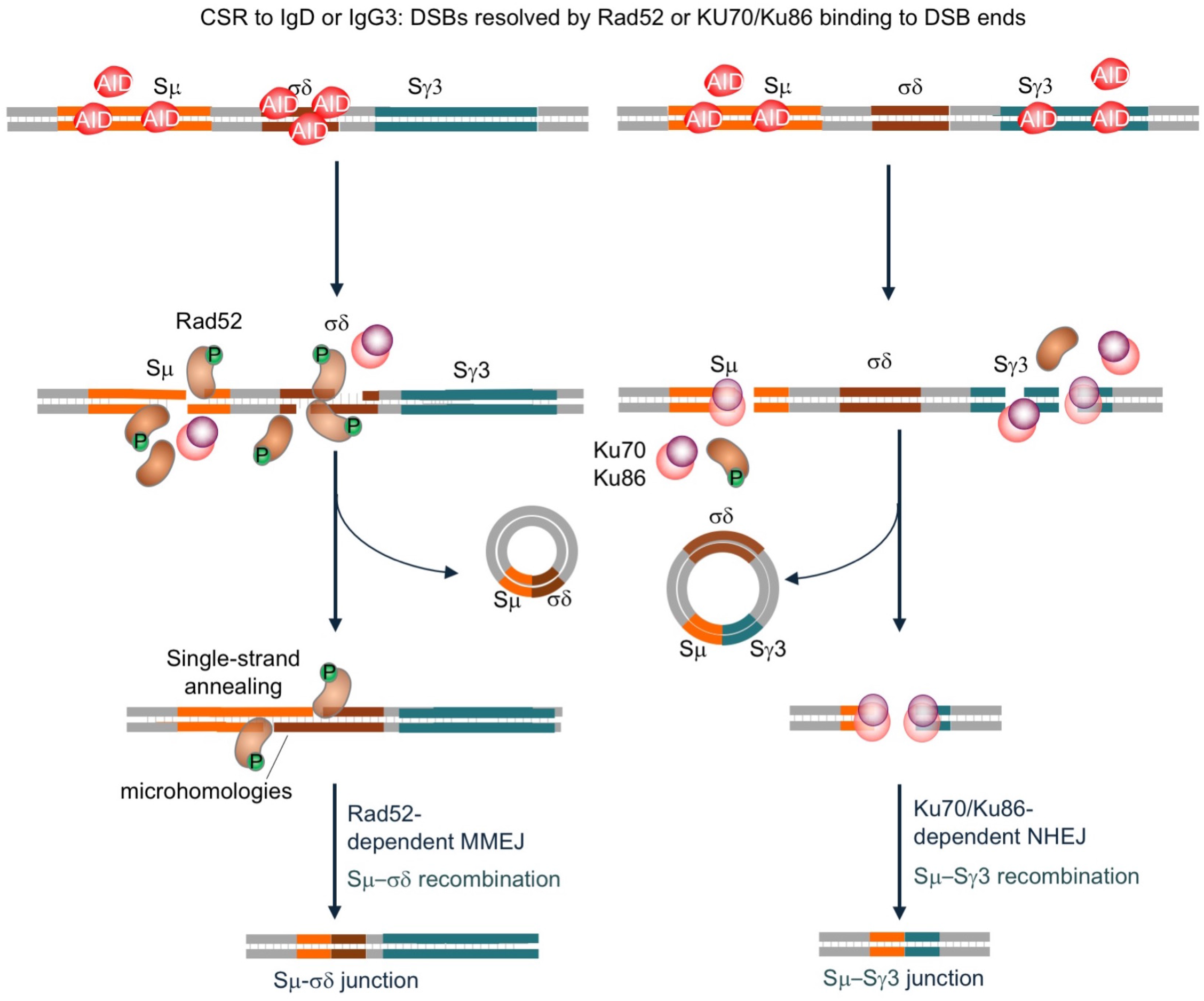
Rad52 mediates Sμ-σδ DNA recombination (CSR to IgD). CSR is initiated by AID-mediated generation of multiple DSBs in the targeted upstream Sμ and downstream Sγ3 (shown here), Sα or Sε regions. Ku70/Ku86, a core NHEJ factor, binds to blunt DSB ends to synapse DSBs by NHEJ involving synapsis and long-range end-joining of S region DSBs, leading to inter-S–S region recombination. This entails deletion of the intervening sequence between S regions as an extrachromosomal circle and leads to CSR to IgG, IgA and IgE. Rad52, an HR element, which is phosphorylated upon CSR induction, binds preferentially to resected DSB single-strand overhangs and facilitates a (Ku70/Ku86-independent) microhomology-mediated A-EJ, which favors intra-S region recombination but can also mediate, particularly in the absence of the NHEJ pathway, inter-S–S CSR. In CSR to IgD, which involves short-range Sμ-σδ recombination. B cells recruit the CSR machinery, including AID, to constitutively transcribed Sμ and σδ regions, and introduce DSBs into these regions. These DSB ends undergo abundant resection, yielding single-strand DNA overhangs. Upstream Sμ and downstream σδ DSB complementary overhangs are rejoined by Rad52 through MMEJ; Upstream Sμ and downstream Sγ3 DSB blunt ends are rejoined by Ku70/Ku86 through NHEJ.

**Supplemental Table 1.**
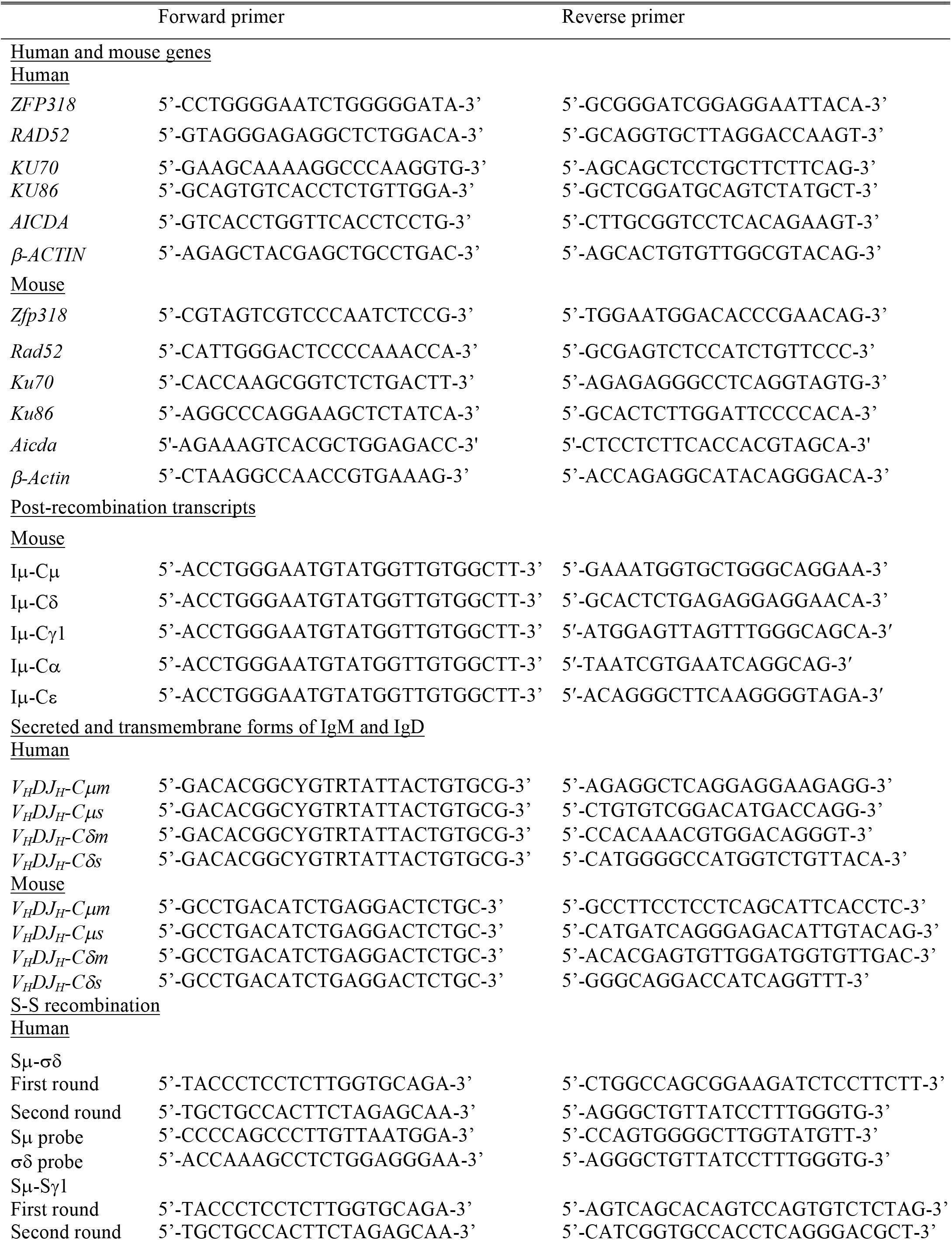
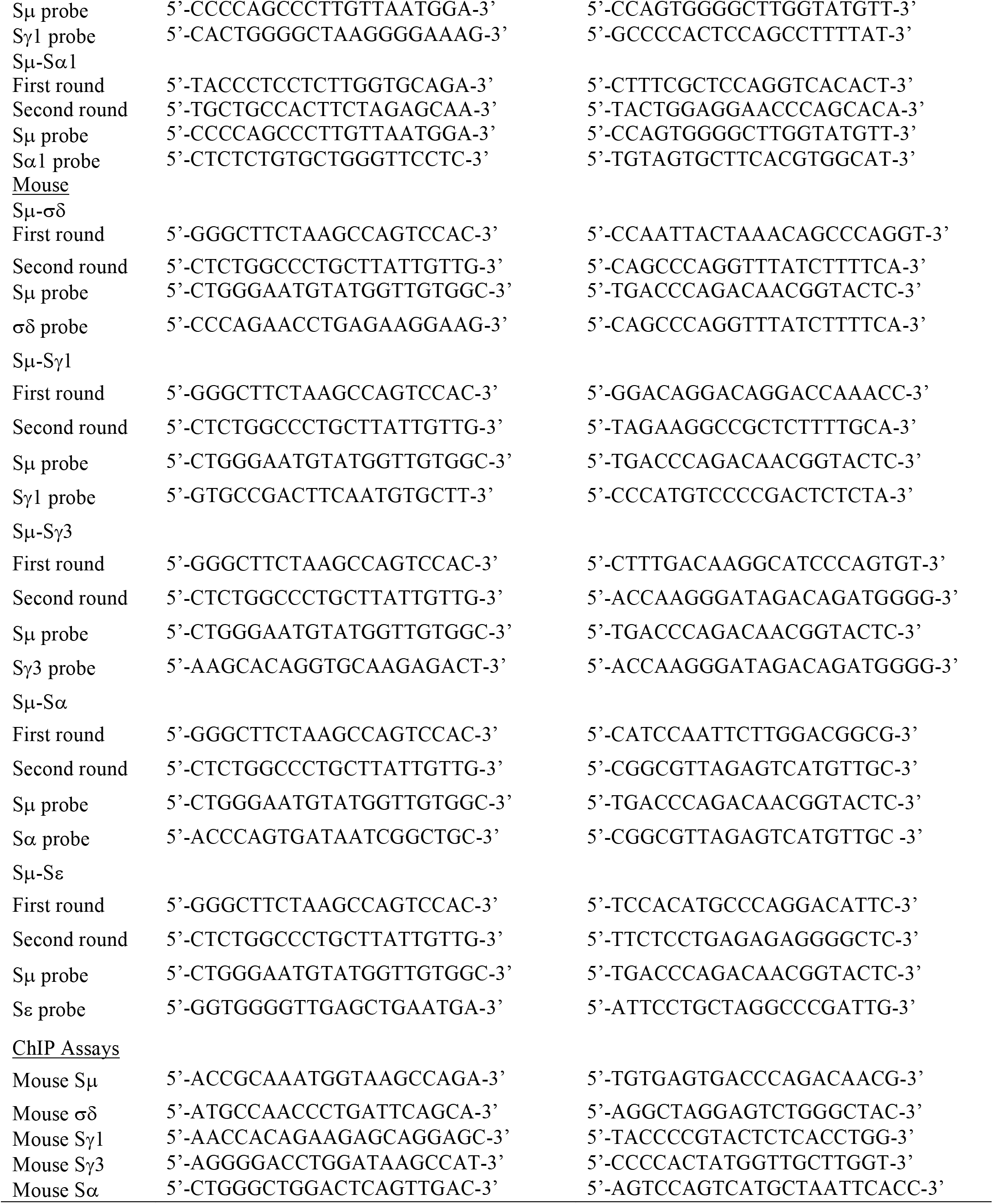
Primers used for this study.

